# 140 Years of mathematical modeling in oncology through AI-assisted curation

**DOI:** 10.64898/2026.01.13.699306

**Authors:** Franco Pradelli, Maximilian Strobl, Sadegh Marzban, François de Kermenguy, Ari Barnett, Katyayni Ganesan, David A. Hormuth, Sara Hamis, Dhananjay Bhaskar, Guillermo Lorenzo, Alexander R. A. Anderson, Jeffrey West

**Affiliations:** Integrated Mathematical Oncology Department, Moffitt Cancer Center, Tampa, FL. USA; Department of Genomic Medicine, Cleveland Clinic Research, Cleveland Clinic, Cleveland, USA; Department of Life Sciences, Imperial College, London, UK; Institute of Cancer Research, London, UK; I-X Centre for AI in Science, Imperial College London, London, UK; Harvard Medical School, Dana-Farber Cancer Institute, Brigham and Women’s Hospital, Boston, MA. USA; Institute of Computational Cancer Biology (ICCB), University Hospital and University of Cologne, Cologne, Germany; Oden Institute for Computational Engineering and Sciences, The University of Texas at Austin, Austin, TX, USA; Division of Systems and Control, Department of Information Technology, Uppsala University, Uppsala, Sweden; Biomedical Engineering Department, University of Wisconsin-Madison, Madison, WI, USA; Group of Numerical Methods in Engineering, Department of Mathematics, and CITEEC, University of A Coruña, A Coruña, Spain

## Abstract

Constructing a comprehensive overview of any scientific field requires accurate literature selection, yet conventional keyword-based searches are susceptible to false positives. This problem is magnified in growing or interdisciplinary fields such as mathematical modeling in oncology that contain a rich but heterogeneous body of literature. Here, a generalizable, context-enriched artificial intelligence pipeline based on large language models (LLMs) curates large scientific corpora according to a user-defined field: mathematical modeling in oncology (>35k publications). Benchmarking against expert evaluation demonstrates high accuracy (ROC AUC∼0.95) and agreement with human judgement (correlation∼0.68), outperforming zero-shot LLM curation Analysis of the curated corpus (∼14k) suggests that Mathematical Oncology s distinct from either Systems Biology and Pharmacokinetics/Pharmacodynamics despite employing overlapping methods. Co-occurring citation network analysis defines nine research clusters focused on a range of applications including drug delivery, optimal control, stochastic modeling, tumor microenvironment, radiation, cancer evolution, and spatial multiscale modeling.

**Significance Statement:** A generalizable, context-enriched artificial intelligence pipeline accurately curates large scientific corpora Analysis of the curated dataset applied to the se of mathematics n oncology provides comprehensive view of mathematical modeling in oncology across 140 years, revealing its shift from fundamental cancer biology towards therapeutic modelling.

## Introduction

Mathematical modeling has shaped oncology since the end of the 19th century, enabling scientists to address fundamental questions such as the role of mutations^2,3^ and therapeutic intervention^4,5^ It has provided both mechanistic^6,7^ and predictive^8,9^ insights into progression and treatment, evolving from initial pioneering investigations into a interdisciplinary, data-driven, and translational scientific landscape^10,11^ Indeed, cancer originates from genetic alterations whose evolutionary trajectories are driven by intricate interactions across molecular, cellular, tissue, and systemic levels, ultimately leading to metastatic dissemination^12^ Each scale offers unique entry points for mathematical exploration, with distinct methods suited to different biological questions.

Among the most established methodologies, deterministic differential equations represent a cornerstone of the field^13^ These are broadly classified into ordinary (ODEs) and partial differential equations (PDEs). ODEs characterize a system through continuous functions n time (e.g. tumor burden, drug concentration, clonal frequency), with classical applications including modeling tumor growth laws^14^ the identification of optimal treatment schedules^15^ and the emergence of resistance^16^ Pharmacokinetics and pharmacodynamics (PK/PD) modeling constitutes the mathematical foundation for characterizing drug–body interactions. Since its emergence in the 1960s^17^ it has become central to drug development, personalized medicine, and therapeutic monitoring^18^ By contrast, PDEs explicitly incorporate spatial dimensions into cancer dynamics^19^ which are often calibrated and validated against preclinical^20^ and clinical^21^ imaging. n parallel (and often in combination) with PDEs, Agent-Based Models (ABMs) capture spatial heterogeneity by simulating individual agents— often cells or molecules—each governed by unique behaviors and interactions^22^ Despite often restricting the analysis to smaller spatial and temporal scales, ABMs excel in capturing spatial heterogeneity^23^ or emergent phenomena^24^ that are difficult to study via differential equations.

The growing focus on heterogeneity and evolutionary dynamics in cancer biology^25^ has further expanded the methodological repertoire to directly address tumor evolution and selection between distinct populations. Probabilistic modelling (e.g. branching and Moran processes)^26^ originally applied to study the effect of mutations^2,3^ currently underpin investigations into tumor plasticity and resistance^27^ While these modeling techniques characterize clonal dynamics and phenotypic switching, evolutionary game theory (EGT) focuses on ecological interactions between different populations^28^ Although EGT applications relatively recent, the framework has proven pivotal for designing therapeutic strategies that mitigate drug resistance and exploit evolutionary trade-offs^28^

Mathematical modeling in oncology has also benefited from integration with machine learning (ML) methods, leveraging the increasing availability of multiscale, multi-modal data. One of the most relevant advancements is the emergence of Mechanistic Learning^29^ which integrates data-driven methods with classical mathematical approaches. Although still in their infancy, these techniques hold promise for clinical translation^30^ and reflect a broader emphasis on clinical applications of mathematical models^31^ A particularly compelling vision is the concept of Digital Twins, whereby patient-specific models—calibrated with longitudinal biological and clinical data—could ultimately forecast therapeutic responses at the individual level^32^

This increasingly complex landscape has been periodically surveyed through reviews and perspectives^10,11,33,34^ Nevertheless, important gaps remain. First, the broader field of mathematical models in oncology (hereafter “mathematical modeling in oncology”) is fragmented across subareas—such as Mathematical Oncology, Systems Biology, and PK/PD—whose boundaries are often unclear. Second, reviews necessarily limit their scope to a few hundred publications, often focusing on specific modeling paradigms or emerging trends. Moreover, although centered o quantitative methodologies, these reviews ironically rarely apply quantitative analyses to their own bibliographic corpora. Bibliometrics, by contrast, provides a systematic and data-driven framework for evaluating the evolution of scientific domains, mapping thematic structures, and uncovering emerging trends across large bodies of literature^35^

The basis of any bibliometric analysis is the selection of a robust publication corpus. Keyword-based searches across scientific databases can provide broad coverage, but often introduce false positives—records that match search terms without being relevant to the analytical question^36^ To address this limitation, we complemented a keyword-based retrieval strategy with automatic curation operated by large language models (LLMs).

Although LLMs have shown strong capabilities in summarizing and interpreting scientific text, hallucinations and other limitations continue to pose concerns on their responsible use in research workflows^37,38^ Therefore, a central aim of this study is to evaluate whether LLMs ca reliably curate bibliographic records for a defined scientific objective.

To this end, we assess nine foundation models from major providers (Claude, Gemini, and OpenAI) and develop a custom architecture that automatically derives contextual evaluation criteria from human-curated scientific publications. We benchmark this approach against expert human evaluation and zero-shot LLM curation, demonstrating high accuracy and agreement with expert judgement (**Fig. 1, 2**).

**Figure 1.**
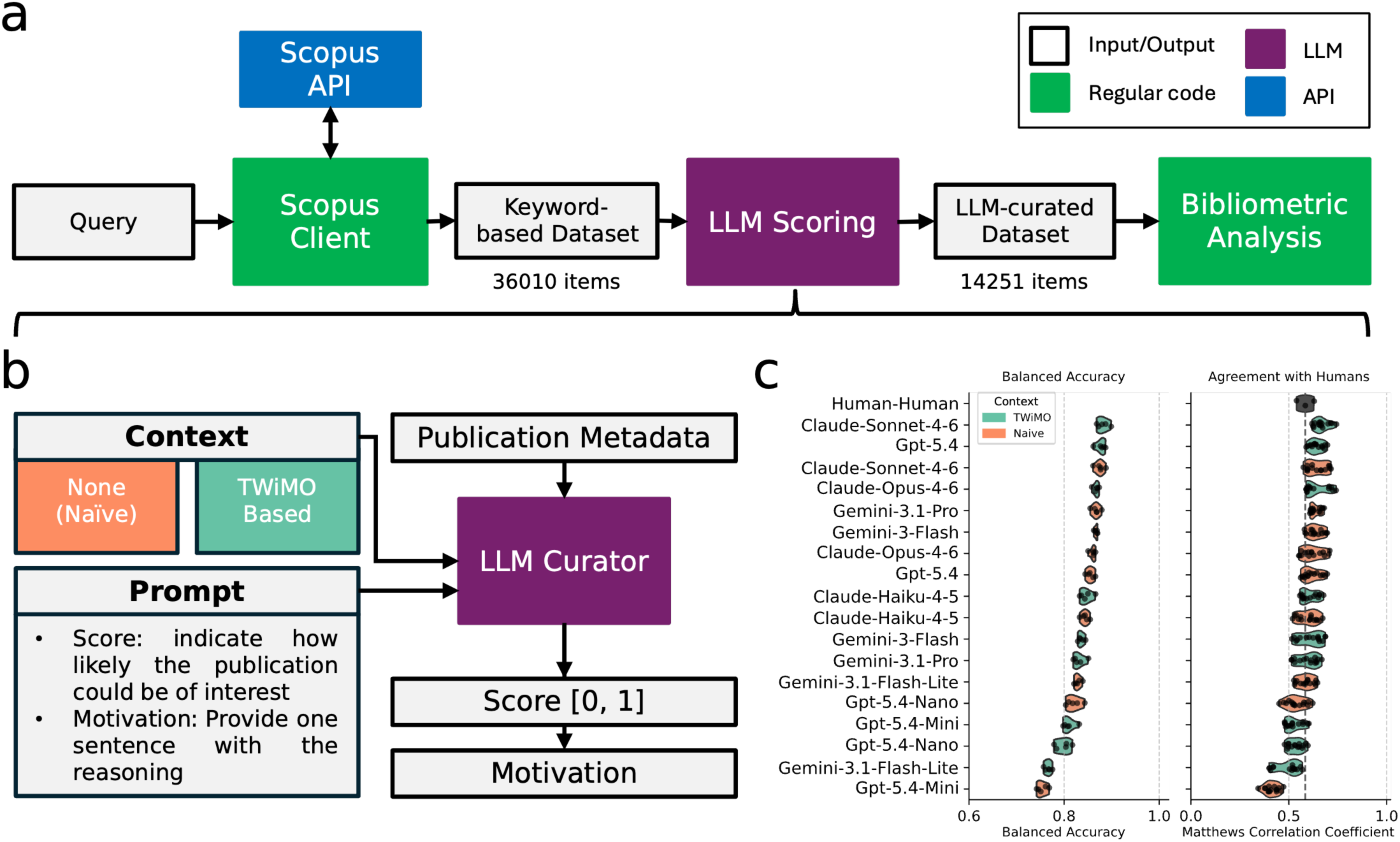
LLM curation effectively evaluate scientific publications. **a)** A schematic of the approach employed in this study to generate an LLM-curated dataset for our bibliometric analysis. **b)** Schematic for the LLM-curation. For each entry of the keyword-based dataset, the LLM-curator was asked to provide a score based on the interest for a mathematical oncologist, and a motivation on its reasoning (an example is provided in the supplementary materials, **LLM Output 3.3**). **c)** Balanced Accuracy and Agreement with human curators (measured with the Matthews Correlation Coefficient) for each foundational model and context strategy. Other evaluation metrics are provided in the supplementary materials (see **Methods**).

**Figure 2.**
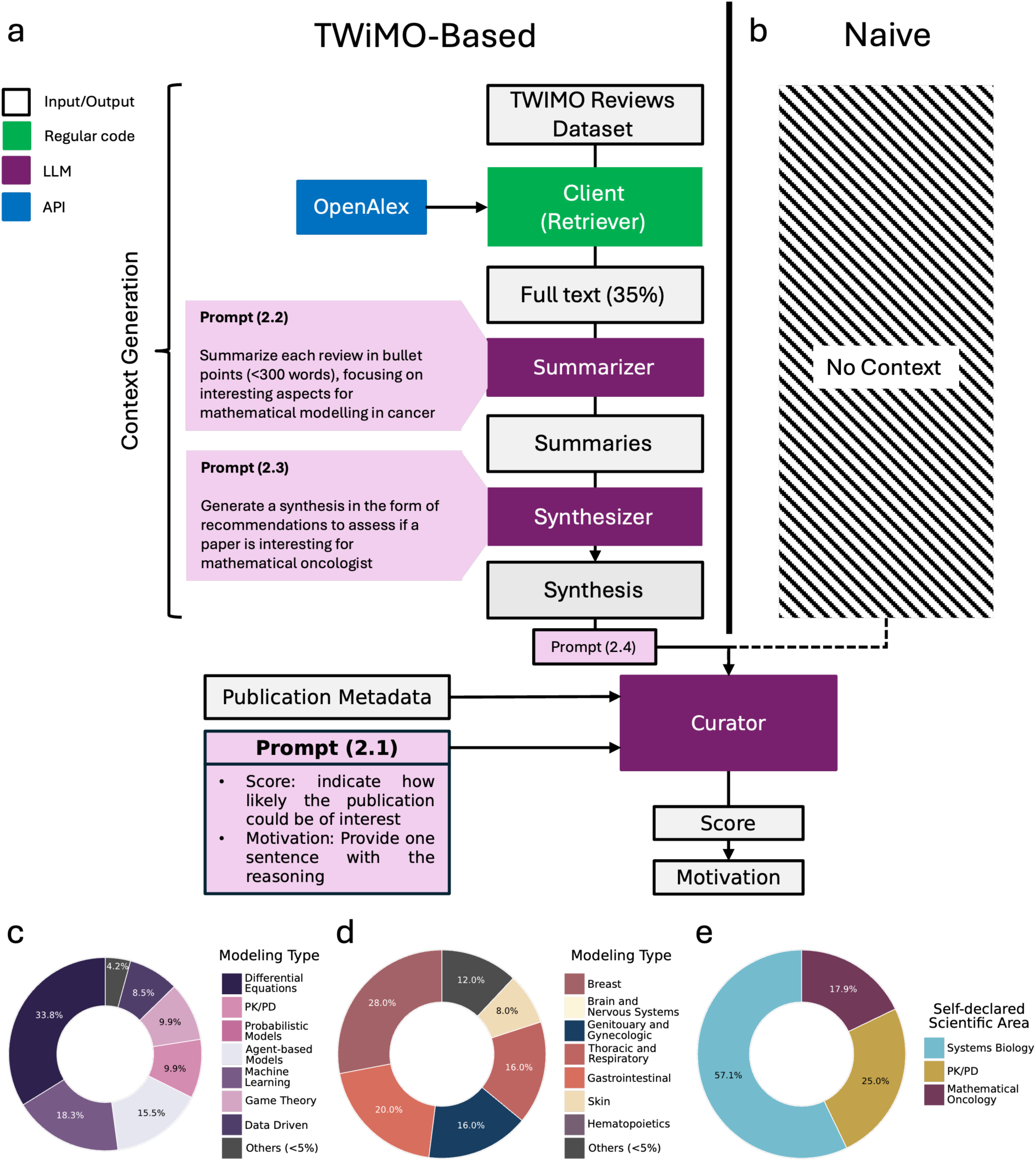
Context strategies to instruct LLM Curators. **a)** The TWiMO-based approach leverages a checklist (Synthesis) automatically generated from the reviews contained in the TWiMO dataset. When possible, we used OpenAlex to download the full text of the review (35% of the cases, 88 reviews). Reviews with no available full text were excluded from the pipeline. Four reviews were associated with an empty PDF in OpenAlex, leading to a total of 84 reviews successfully employed for the context strategy. **b)** The Naïve strategy provides no context to the LLM Curator. **c)** Modeling methods, **d)** Cancer Types, and **e)** Scientific Areas represented in the dataset used for the TWiMO-based context. The prompts indicated in the image are meant to explain the meaning of the instructions and do not correspond to the exact text employed in each situation. For the exact prompts, see **Supplementary Appendix 2.**

This approach allowed us to combine inclusiveness with precision. From an initial keyword-based dataset of more than 35k publications, the LLM-curation pipeline identified approximately 14k records relevant to mathematical modeling in oncology (LLM-curated dataset). We use this curated corpus, together with domain expertise and bibliometric analyses, to provide comprehensive perspective the role of mathematics in research and clinical management, with particular emphasis on the emergence of Mathematical Oncology.

Analyzing the LLM-curated dataset, we demonstrate how mathematical modeling in oncology expanded as a translational, impactful, and multidisciplinary field (**Fig. 3** and **4**). We identify the dominant themes that have shaped the discipline and delineate Mathematical Oncology as distinct from two other closely related fields—Systems Biology and PK/PD (**Fig. 5**). We examine the research interests of influential and prolific contributors and, through bibliographic coupling, reveal distinct research lines shaping the current scientific landscape and its temporal evolution (**Fig. 6**). We show that key drivers of the field’s evolution include, but are not limited to, the integration of spatial modeling, imaging, evolutionary and ecological dynamics, stochastic modeling, drug delivery, optimal control, and multiscale modeling (**Fig. 6**). Collectively, our analysis highlights an ongoing conceptual shift in mathematical modeling in oncology, from fundamental cancer biology toward translational clinical applications. This evolution has placed increasing emphasis on therapeutic modeling and its relevance for clinical decision-making, coinciding with the rise in prominence of Mathematical Oncology. In conclusion, our findings indicate that Mathematical Oncology is more than the application of mathematics to cancer research. We propose to define it as the use of interpretable mathematical models—integrating clinical, biological and physical knowledge— to enhance cancer screening, understand disease evolution, guide therapy, and strengthen forecasting.

**Figure 3.**
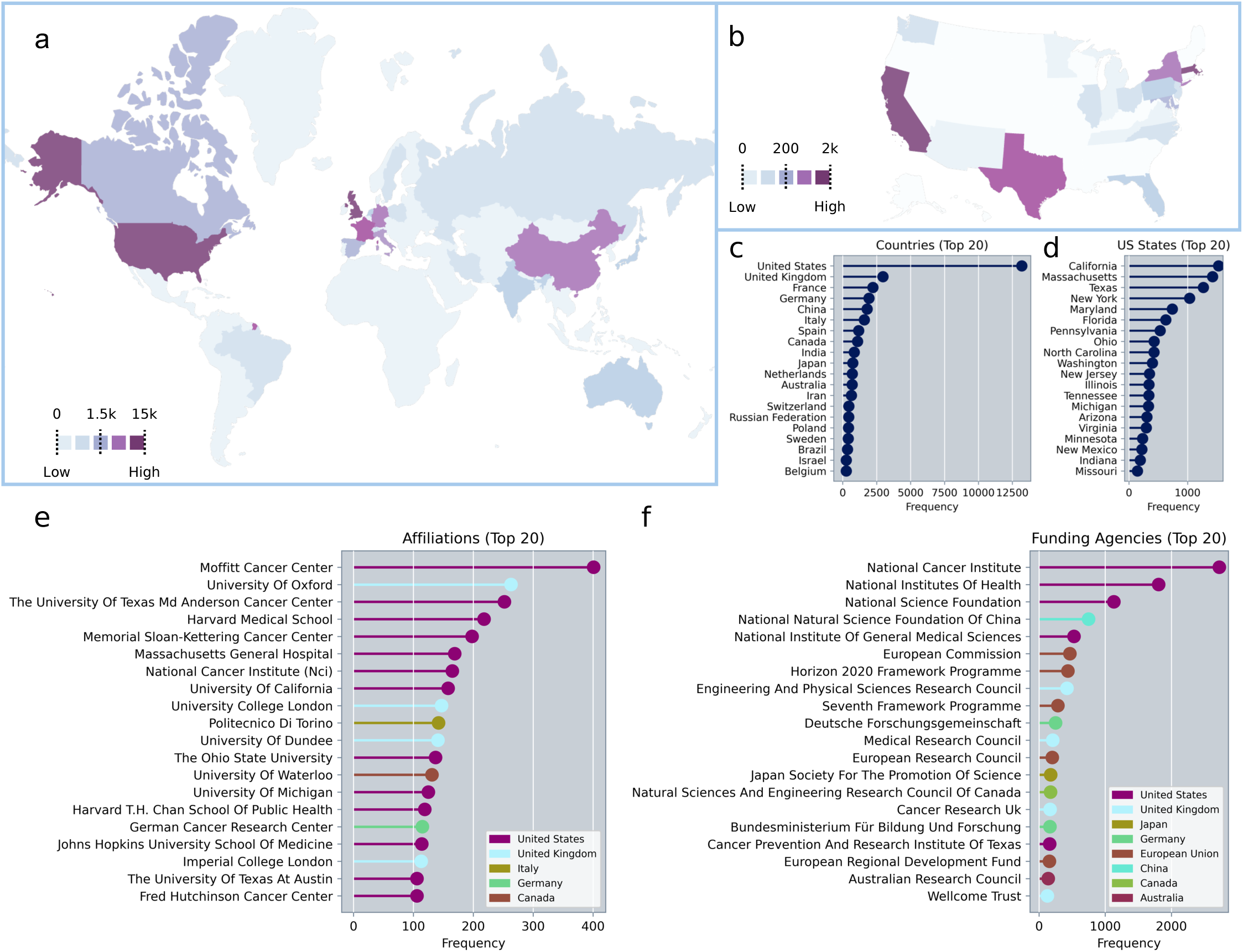
Geographic distribution of mathematical research in oncology (LLM-curated dataset). **a)** Heatmap showing the number of documents of each country and **b)** of each state in the U.S (the scale is logarithmic). **c)** Number of documents per country and **d)** each state in the U.S. Despite the prominence of western countries, the distribution shows relevant contributions from Asia (China, India, Japan, Pakistan), Middle-East (Iran, Saudi Arabia), and South America (Brazil). **e)** Most represented institutions in the LLM-curated dataset, colored for their country. The U.S. is the single most represented country with 13 institutions, including major cancer centers such as Moffitt, MD Anderson, Memorial Sloan-Kettering, and The National Cancer Institute. The second are the U.K., with four institutions: Oxford, University College London, Dundee, and Imperial College. **f)** Most represented funding sponsors in the LLM-curated dataset, colored for their country. They are mostly located in western countries (five in the U.S., four in the U.K., and European Union), but Chinese and Japanese funding agency occupy high positions in the ranking.

**Figure 4.**
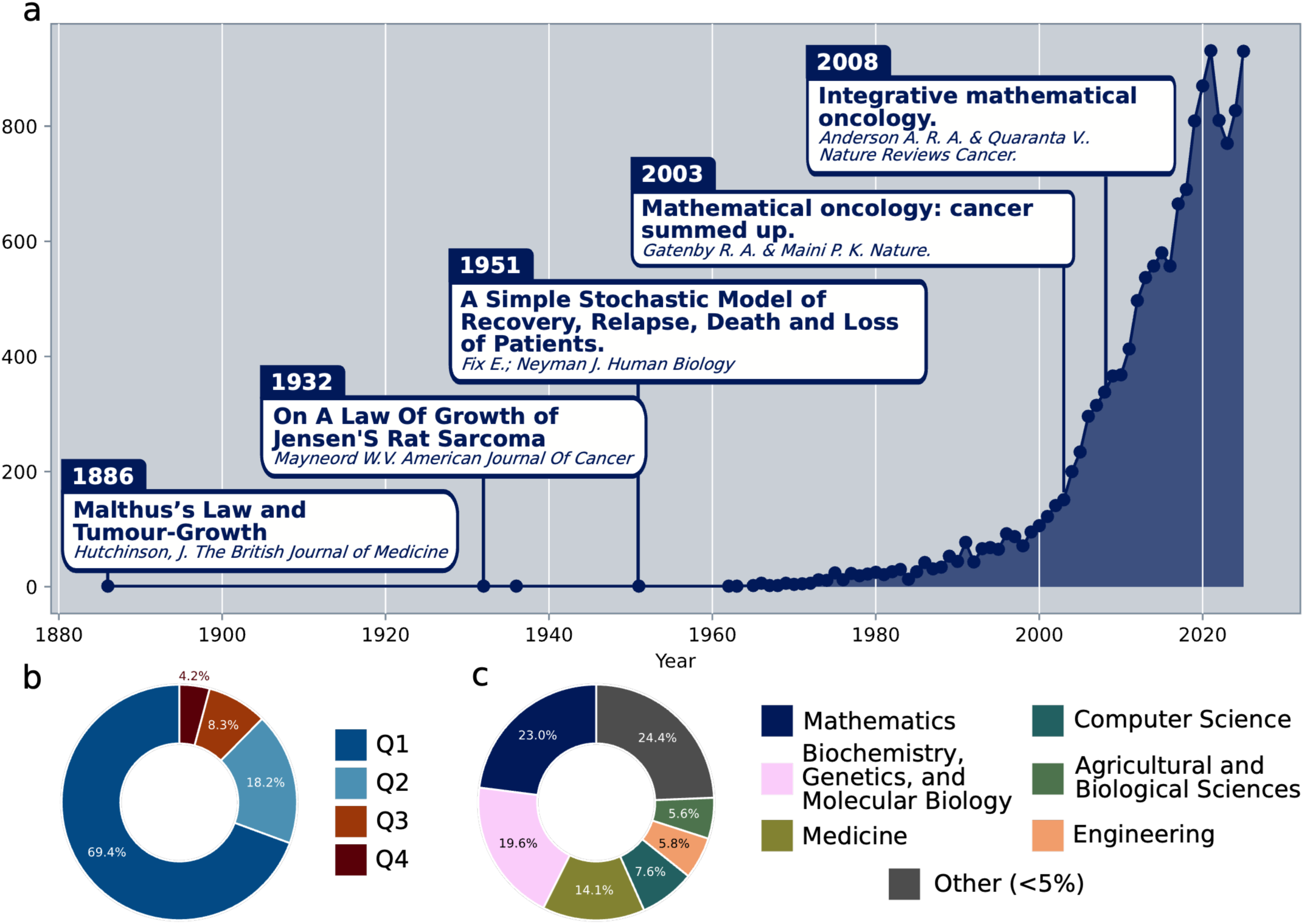
The expansion and impact of mathematical research in oncology. **a)** Number of documents published in the area in time; field-defining publications are highlighted, including the first entry in the dataset (1886), the first publications integrating biological data (1932) and clinical data (1951) with mathematical modeling, the first paper mentioning “Mathematical Oncology” (2003), and the paper that served as a manifesto of the field (2008). **b)** Journal quartile for each publication in the llm-curated dataset. **c)** Journal subject areas for each publication in the query-based dataset. The group “Other” includes all ASJC areas representing less than 5% of the publications in the query-based dataset.

**Figure 5.**
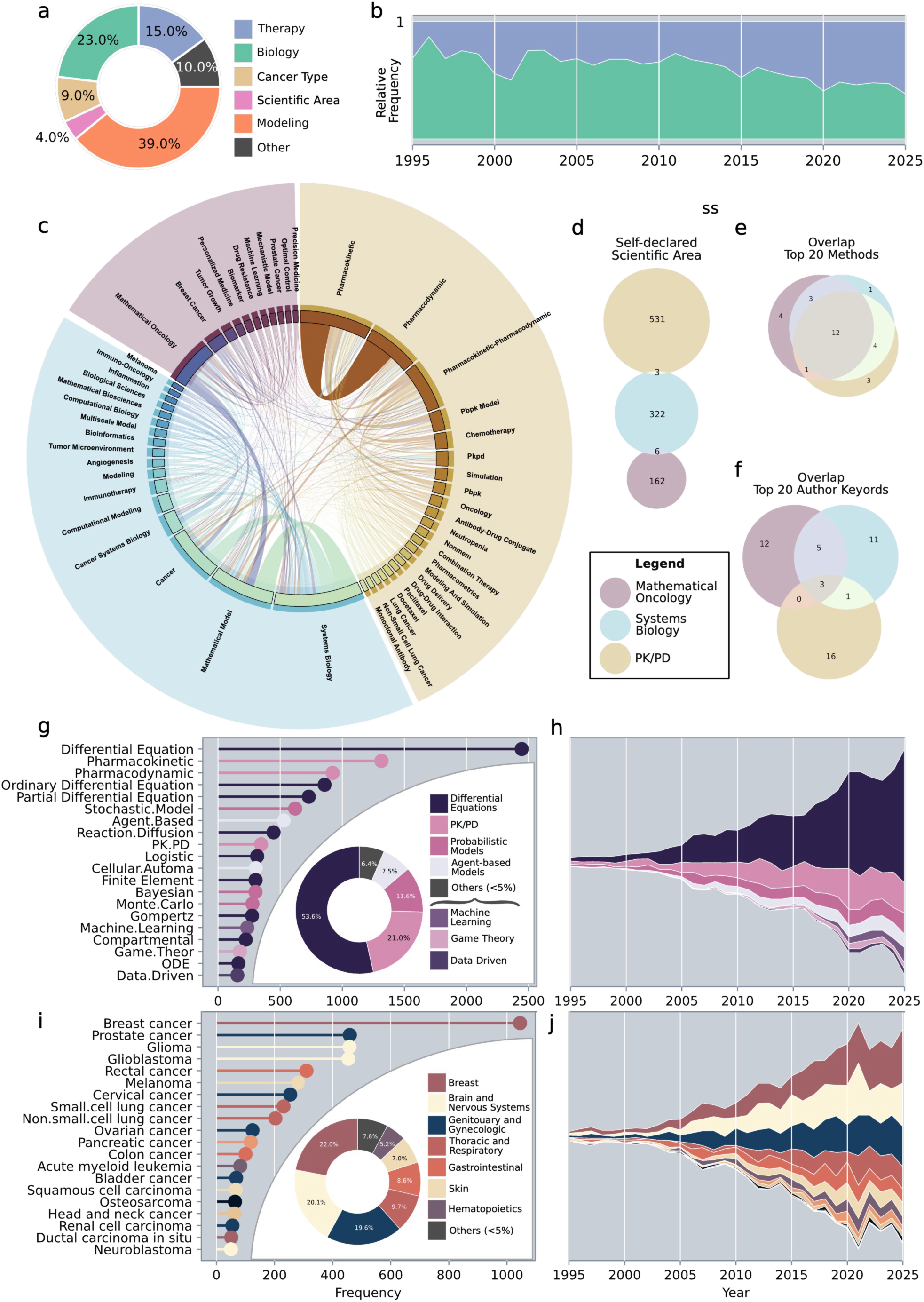
Primary research topics and areas in the llm-curated dataset. **a)** Semantic group distribution of the top 100 AKs (see full list in **Fig. S6a**). **b)** Relative frequency (normalized to 1) of the AKs associated to therapeutic applications, and to cancer biology (full list in **Fig. S6b-S6c**). **c)** Co-occurrence network between the most popular AKs in each scientific area (Mathematical Oncology, Systems Biology, PK/PD). The color of the outer circle represents the scientific area of each AK (also evidenced in shade). The width of each chord in the chart is proportional to the co-occurrence between each pair of terms (see **Methods**). **d)** Overlap between the scientific areas declared in the AKs of the document self-identifying with one. **e)** Overlap between the top 20 methods for the documents self-identifying in each scientific area. **f)** Overlap between the top 20 AKs for the documents self-identifying in each scientific area. **g)** 20 most represented regexes associated with specific modeling method, and **h)** major modeling categories in the last 30 years (1995-2025). The coloring in **g)** and **h)** reflects the major modeling category, and the pie chart shows the distribution of each major modeling category in the dataset (legend next to the pie chart). **i)** The 20 most represented regexes associated with specific cancer type, and **j)** representation of cancer types (by location) in the last 30 years (1994-2024). The coloring in **i)** and **j)** reflects the tumor location and the pie chart shows the distribution of each location in the dataset (legend next to the pie chart).

**Figure 6.**
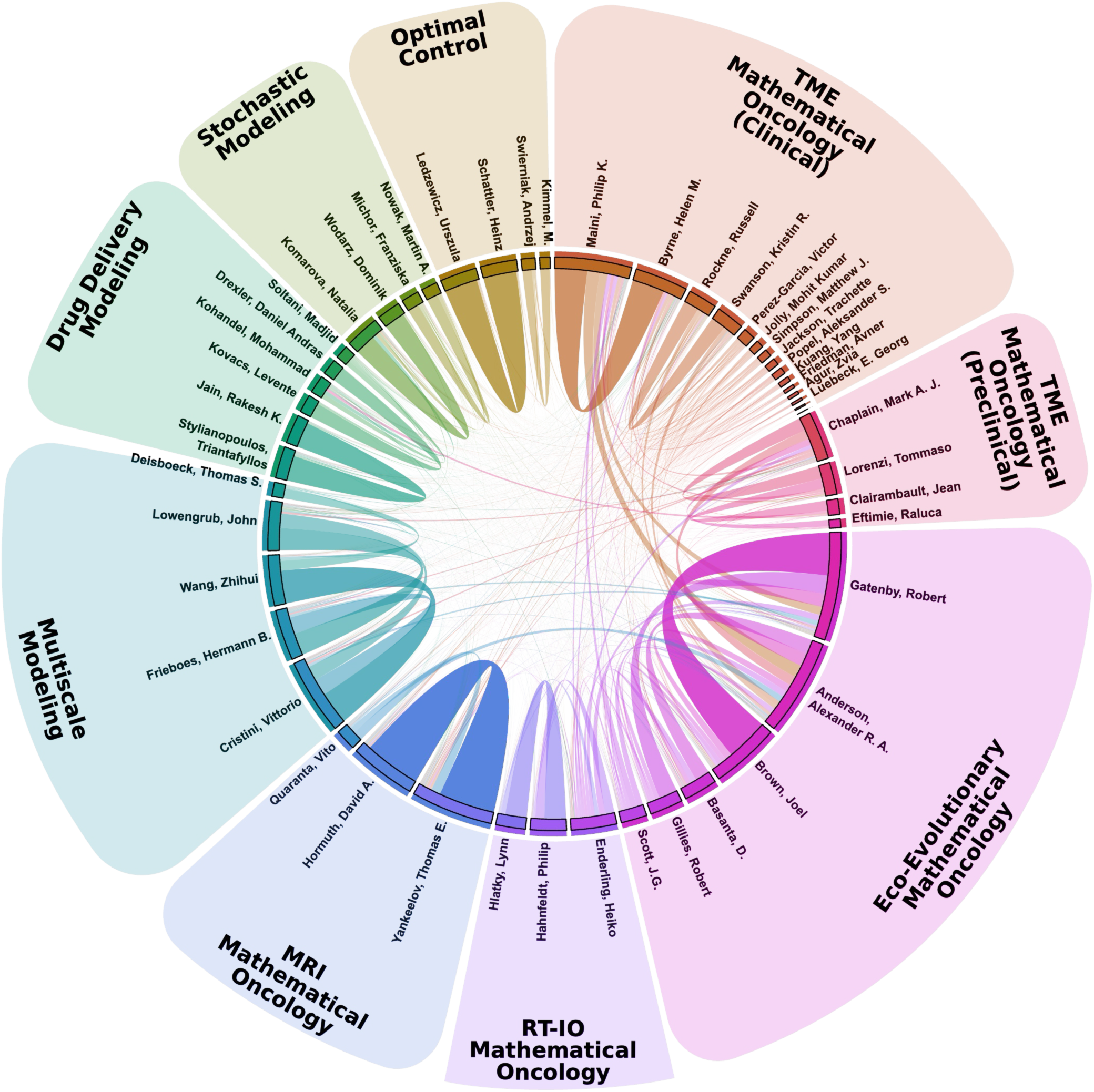
Bibliographic coupling between prominent contributors to mathematical research in oncology. The width of each chord is proportional to the tendency for each couple of authors to cite the same references (so, to work on the same concepts; see **Methods**). The color of the outer circle corresponds to the cluster of each author (9 clusters in total, also evidenced in the shade). The name of each author cluster is based on the frequent terminology adopted within the cluster. If the cluster adopts frequently the term “Mathematical Oncology”, we included this term in the name of the cluster. To improve readably, Kim, Yangin and Kim, Jane (Cluster “TME Mathematical Oncology (Clinical)”) have been hidden from the visualization.

## Results

### Large language models equal human experts in publication curation

A known limitation of keyword-based bibliographic retrieval is the inclusion of false positives, i.e. documents that match the query but fall outside the scope of interest. This issue arises from the inherent ambiguity and flexibility of language (e.g., false-positives via homonyms) and is present in our keyword-based dataset. Manual evaluation of 200 randomly sampled documents by three independent experts indicated only 66% were deemed relevant by majority vote, whereas 30% were unanimously classified as irrelevant (**Fig. S1, Methods**).

Although manual curation could, in principle, resolve this issue, the scale of the dataset (36,010 documents) makes such an approach impractical. We therefore developed and assessed a novel large language model (LLM) pipeline to perform the task of bibliometric evaluation and curation.

Our LLM pipeline achieves comparable performance to expert evaluators in identifying publications pertinent of interest for mathematical oncology (**Fig. 1**). The core of our workflow is scoring step (**Fig. 1a**), in which an LLM curator assigns a relevance score between 0 and to each document, and provides a brief justification on its reasoning (**Fig. 1b, Methods**). The best-performing model (Claude Sonnet 4.6) attains a balanced accuracy of 0.88 and a mean agreement with human evaluators of 0.67 (**Fig. 1c**), exceeding the nter-annotator agreement among human experts (0.58).

Providing additional contextua information generally improves the performance of LLM curations, although the effect varies across foundational models. We evaluated three models (Claude, Gemini-3.1, GPT-5.4) in three different model sizes. For each, we compared two strategies: naïve approach without contextual guidance, and context-enriched approach based on evaluation criteria automatically generated from the publications featured o “This Week in Mathematical Oncology” (TWiMO, see **Methods, Fig. 2**). The TWiMO-based strategy improved performance across all Claude models and most GPT variants, but consistently reduced performance for Gemini models. These results are consistent with known trade-offs in context utilization across LLM architectures, whereby increased context ca enhance or impair performance depending on model design and capacity^39^

Applying the LLM-based filtering step resulted in a refined dataset of 14,251 publications, a reduction of 60.4% of manuscripts from the keyword-based dataset, which forms the basis of subsequent analyses (LLM-curated dataset).

### LLM-curated dataset reveals the international and translational scope of mathematical modeling in oncology

Analysis of the institutions reported in the LLM-curated dataset reveals the global scope of mathematical modeling in oncology (**Fig. 3**). The data demonstrate a marked predominance of the United States (**Fig. 3a–d**): of the 20 leading institutions (**Fig 3e**), 13 are U.S.-based, including major cancer centers such as Moffitt, MD Anderson, Memorial Sloan-Kettering, and The National Cancer nstitute. Within the U.S., California, Massachusetts, and Texas ead in publication output. Although Florida hosts the single most prolific institution (Moffitt), the cumulative contributions of multiple institutions make other states more productive overall. The United Kingdom ranks second, with four institutions, followed by institutions from Italy (Politecnico di Torino), Germany (German Cancer Research Center), and Canada (University of Waterloo).

Historically, both academic and clinical institutions have contributed to the expansion of mathematics in oncology. The analysis of the institutional ranking across different time frames (**Fig. S2**) demonstrates that hospital and cancer centers are always represented in the apical position, demonstrating a strong translational nature of the discipline.

The analysis of the funding data (**Fig. 3f**) suggests a link between institutional strength and to dedicated research funding. All the clinically oriented sponsors (e.g. NCI, NIH, Medical Research Council) are concentrated in the U.S. and the U.K., which lead the institutional ranking (**Fig. 3e**). The U.S. and the European Union stand out as the primary sponsors, with five and four major agencies respectively. The National Natural Science Foundation of China also ranks prominently, reflecting increasing national investment. Although relatively few individual institutions from non-Western countries appear among the top contributors, **Fig. 3c** highlights substantial activity across Asia, the Middle East, and South America, attesting to the increasingly global diffusion of the field. This trend is further corroborated by temporal analyses (**Fig. S2**), which depict a clear expansion of mathematical approaches to cancer research over time.

### Mathematical modeling in oncology reveals continued expansion and interdisciplinary impact

The geographic expansion of mathematical modeling in oncology paralleled by marked increase in publication output (**Fig. 4a**). Early links between mathematics and cancer biology can be traced back more than a century however, sustained growth emerged only in the late 1990s, together with the rise of computational power. This expansion supported the inception of Mathematical Oncology as an independent field. The first publication explicitly mentioning this term dates to 2003^40^ and precisely delineated by Anderson & Quaranta in the foundational publication “Integrative Mathematical Oncology”^11^

While a large share of publications n the area appear in journals within computational and mathematical biology, a substantial fraction are published in multidisciplinary (e.g. *PNAS*) or clinical oncology (e.g. *Cancer Research*) journals, underscoring the translational and interdisciplinary focus of the field (**Fig. S4a**).

Analysis of Journal quartiles further demonstrates that expansion is paralleled by consistent quality (**Fig. 4c**). Fourteen of the twenty most frequent journals (including all of the top ten) are classified in the first quartile of their area (Q1), and 69% of all articles in the LLM-curated dataset appear in Q1 journals (i.e. the most cited journals in their relative area; see **Methods**). This proportion is striking when compared to independent reports estimating the average Q1 publication rate at approximately 30%^41^ highlighting the wide-spread interest and impact of Mathematical Oncology research.

The interdisciplinary scope of the field is further reflected by the distribution of the journals’ subject areas (**Fig. 4d**). Each journal in Scopus is classified in one of more scientific areas following the All Science Journal Classification (ASJC), allowing us to map each document to its scientific areas (ASJC areas see **Methods**). Publications are primarily classified under “Mathematics,” followed by “Biochemistry, Genetics, and Molecular Biology,” “Medicine,” “Computer Science,” “Engineering,” and “Agricultural and Biological Sciences” illustrating how mathematical modeling in oncology bridges computational and mathematical methodologies with biological and clinica sciences. This cross-disciplinary role has remained consistent over time (**Fig. S3e,h,k,n,q**), with more than 50% of publications persistently clustered in “Medicine,” “Biochemistry, Genetics, and Molecular Biology,” and “Mathematics.” Longstanding venues such *Journal of Theoretical Biology* and *The Bulletin of Mathematical Biology* have played centra roles in mathematical modeling in oncology, while more recent journals—including *PLoS ONE Scientific Reports*, and *PLoS Computational Biology*—have expanded the field’s reach.

Open-Access publications and preprints also appear to impact the field’s influence. Citation analysis of the LLM-curated dataset reveals average of 32.23 citations per article (median: 12; **Fig. S4a**), with an average annual rate of 3.04 (median: 1.44; **Fig. S4d**). Open Access (OA) publications exhibit higher citation counts overall (**Fig. S4b, S4e**), with significant differences observed among the four OA modalities considered in Scopus (Gold OA, Hybrid Gold OA, Bronze OA, and Green OA; see **Methods**). Journals offering temporary OA tend to achieve the highest total citation numbers, but availability in public repository also increases publication impact (**Fig. S4c, S4f,** and **Table S1**). These findings indicate that unrestricted accessibility—especially temporary or via repositories—consistently enhances the visibility of research.

### A shift from ancer biology to therapy modeling traces the rise of Mathematical Oncology

The strong impact and expansion of mathematical modeling in oncology must rely compelling research investigations. To have further insights into the primary topics in the field, we analyzed the usage of the AKs (**Fig. 5a-d, 5f, Fig. S5**), and the frequency of terms (see **Methods**) related to mathematical modeling techniques (**Fig. 5g-h**) and cancer types (**Fig. 5i-j**).

The interdisciplinary orientation of the field is reflected by the choice of the author keywords, which divide almost equally between modeling-related terms, and concepts related to cancer types, biology, and therapy (**Fig. 5a, Fig. S5a,** see **Methods**). Commonly used author keywords include generic terms as *“Mathematical Model”* and *“Cancer”* but also common therapeutic modalities (e.g. *“Chemotherapy” “Immunotherapy”*), modeling techniques (e.g. *“Agent Based Model”*), cancer types (e.g. *Breast Cancer”*), and terminology connected to specific aspects of cancer biology (e.g. *“Tumor Growth”*, “*Angiogenesis*”).

Despite the prevalence of terms related to cancer biology (**Fig. 5a**), temporal trends reveal a clear shift from cancer biology toward therapy (**Fig. 5b**). While usage of terms such as *Metastasis” Cell Cycle*”, *Hypoxia*”, and others has declined in the past ten years (**Fig. S5b**), terms linked to therapy have increased (**Fig. S5c**), reflecting growing interest in treatment optimization and data-driven approaches (as also noted by others^29^).

This conceptual transition toward therapy-related seems to be closely tied to the rise of Mathematical Oncology as a distinct subfield Frequently, authors include AKs related to their scientific area, the most prominent being “Systems Biology”, “Mathematical Oncology”, and “PK/PD” (**Fig. S5d**) Among these terms, *“Mathematical Oncology”* has see the most important and consistent increase in the last 10 years (**Fig. S5d**). Thus, we analyzed the use of AKs between documents identifying themselves as part of the subareas “Systems Biology”, “Mathematical Oncology”, and “PK/PD Modeling” (see **Methods**).

The co-occurrence network of AKs (**Methods**) demonstrates the three areas of mathematical modeling in oncology differentiate for different interests and angles on cancer research (**Fig. 5c**). Mathematical Oncology shows a strong connection with translational applications, such as precision medicine, personalized medicine, optimal control, and drug resistance. On the other side, Systems Biology shows specific interest for fundamenta aspects of biology (e.g. angiogenesis, tumor-immune interactions), while PK/PD demonstrate a strong focus o drug delivery (especially for chemotherapy).

Interestingly, authors demonstrate a strong tendency to identify their work within only one of the three scientific (**Fig. 5d**), reinforcing the formation of strong field identity. This identity s not constructed o the modeling techniques of choice, but rather on the scientific question, as demonstrated by the overlap between common modeling techniques (**Fig. 5e**) and the separation between common AKs (**Fig. 5f**).

The analysis of the AKs related to modeling techniques and cancer types clearly shows that some methods and diseases are more studied than others (**Fig. S5a**). To complement the AKs analysis, we performed text mining of titles, abstracts, and keywords, quantifying occurrences of curated set of regexes related to modeling approaches or cancer types (see **Methods**). This allowed s to investigate these two semantic spaces regardless of the author biases, and the other limitations related to AK choice (e.g. space, missing values). Regexes enable a more precise identification of specific character patterns, allowing flexible matching (for example, the regex “*X.ray*” matches both “*X ray*” and “*X-ray*”).

The analysis of modeling approaches highlights the historical predominance of differential equations, pharmacokinetic/pharmacodynamic models, and probabilistic approaches (**Fig. 5d–e**). Coherently with the established relevance of differential equations in the field, this group is the most represented in the last 30 years (**Fig. 5d**). Of note, terms related to classic differential equations such as *Logistic*”, *Compartmental*”, and *Gompertz*”, appear in the top 20. Over the past three decades, ABMs, EGT, and ML have emerged strongly, with regexes such as *Agent.Based*,” *Cellular.Automata*,” *Game Theory, Machine Learning,* and *Artificial Intelligence* appearing among the 20 most frequent terms, reflecting the rise of alternative approaches, and data-driven paradigms.

Tumor-type analysis demonstrates the prominence of breast, prostate, and brain cancers, particularly gliomas and glioblastomas, as the most frequently studied tumors (**Fig. 5f–g**), accounting for over half of all publications in the past 30 years. This focus is likely motivated by the high incidence of breast and prostate cancer^42^ and the marked aggressiveness and poor prognosis of gliomas and glioblastomas^43^ which have historically motivated substantial research efforts. Additional focus areas include rectal cancer, melanoma, cervical cancer, and lung cancers, while head and neck and endocrine tumors remain comparatively underexplored (compared to their respective cancer incidence^42^), representing opportunities for future research.

Regex co-occurrence analysis provides further insight into methodological development. The co-occurrence patterns of tumor-related and modelling-related regexes suggest that cancers are typically modeled first with differential equations PK/PD, and subsequently with additional methods, like ABMs, game theory, and machine learning (**Fig. S6**). This trajectory suggests that the evolution of mathematical modeling in oncology is structured less around individual methodologies than around tumor-specific research programs, in which increasing biological insight and data availability enable and require progressively more sophisticated modeling frameworks

### Bibliographic coupling reveals the scientific landscape of mathematical modeling in oncology

While the analysis of the terminology reveals how researchers frame their ow work, examining the relationship between leading contributors shows how distinct groups have shaped the different research lines in mathematical modeling in oncology. To map these relationships, we used bibliographic coupling rather than co-authorship, which be confounded by authorship practices and may not reflect actual intellectual proximity^35^ We therefore constructed a chord diagram representing the bibliographic coupling among the top 50 authors, where link thickness indicates the degree of shared citations, providing an unbiased measure of vicinity in scientific interests (**Fig. 6** see **Methods**) Clustering by connectivity similarity revealed nine groups with distinct research themes and terminologies (hereafter, author clusters), reflected in their of AKs, their preferred mathematical modeling method, and their primary oncologic focus (**Fig. S7**).

A major research trajectory applies partial differential equations and ABMs to spatial interactions and pattern formation in cancer supporting sustained efforts to model the tumour microenvironment, including angiogenesis and spatial heterogeneity. This tradition is captured by the “TME Mathematical Oncology (Clinical)” and “TME Mathematical Oncology (Preclinical)” clusters^33^ the former emphasizes clinically oriented applications, particularly brain tumours and immunotherapy, whereas the latter focuses on biological mechanisms such as chemotaxis and haptotaxis (**Fig. S7a,b**). These clusters include early contributors to the field, including P.K. Maini, H.M. Byrne, and M.A.J. Chaplain (**Fig. S8b**). Notably, Maini and Gatenby co-authored the first paper explicitly introducing the term “Mathematical Oncology”^40^

The recognition that cancer progression is shaped by spatial and ecological interactions has fostered a strong interface between oncology, ecology and evolutionary theory. This perspective defines the “Eco-Evolutionary Mathematical Oncology” cluster, which focuses on ecological and evolutionary models of tumour growth and treatment response (**Fig. S7c**). The authors in this cluster have made seminal advances linking evolutionary dynamics to clinical outcomes, especially using agent-based models and game theory (**Fig. S7c**). A major translational outcome of this work is adaptive therapy (AT)^44^ which leverages tumor heterogeneity and cell–cell competition to control disease progression through flexible treatment schedules. Mathematical modeling has been pivotal in evaluating these schedules, which have shown encouraging results in preclinical studies across multiple tumor types and proved successful for metastatic castrate-resistant prostate cancer^34^

Radiotherapy exemplifies the long-standing influence of mathematics and physics oncology, with classical models such as the linear-quadratic providing the foundation of clinical practice. Recent work extends this foundation by integrating radiation-response models with tumour dynamics, DNA damage and immunotherapy (**Fig. S7d**). The “RT-IO Mathematical Oncology” cluster captures this trajectory, including recent efforts to use mathematical modeling to develop personalized radiotherapy schedules^45^

Medica imaging provides a unique window on spatial tumour development and an important source of quantitative data for mathematical modelling. The “MRI Mathematical Oncology” cluster integrates medical imaging (especially magnetic resonance imaging, or MRI) with mathematical models (**Fig. S7e**). Authors in this group recognized for linking imaging-derived biomarkers with mechanistic and predictive models of tumour growth, with representative applications in breast and brain cancers^46^

Pharmacokinetic and pharmacodynamic modelling have a central role in drug development. An active area of research seeks to connect PK/PD frameworks with spatially and biologically informed models of tumour progression and drug response. The “Multiscale Modeling in Cancer” cluster focuses on multiscale models of tumour progression and their integration with machine learning for predictive medicine bridging systemic, tissue and cellular scales through PK/PD and agent-based modelling^47^ (**Fig. S7f**). The “Drug Delivery Modeling” cluster emphasizes PK/PD and machine-learning approaches to simulate drug distribution within the tumour microenvironment^48^ (**Fig. S7g**).

Somatic mutations are core drivers of carcinogenesis and the evolution of resistance. Given the randomic nature of such events, probabilistic models have been used to estimate the likelihood of carcinogenic events, somatic evolution and treatment response since the mid-twentieth century^2^ This perspective is represented by the “Stochastic Modeling” cluster, which distinguishes itself for seminal efforts to apply probabilistic models to cancer development and therapeutic resistance^49^ (**Fig. S7h**).

Therapeutic decision-making in oncology often requires balancing benefits against risks. This trade-off has motivated extensive work on treatment optimization across many research lines. Optimal control theory provides a natural framework for this challenge, and has been deeply investigated in the “Optimal Control” author cluster, with applications ranging from chemotherapy dosing, to tumor-immune interactions^9^ (**Fig. S7i**).

Across these groups, Mathematical Oncology emerges as the broadest disciplinary label for mathematical models of cancer It appears across multiple clusters, whereas no alternative field-level term shows comparable reach (**Fig. S7**). Clusters that do not se “Mathematical Oncology” generally rely on broader expressions such as “mathematical modelling” or method-specific descriptors such as “stochastic modelling” and “multiscale modelling”, which do not convey a distinct oncological identity.

The temporal evolution of bibliographic coupling evidence the increasing relevance of spatial modeling in the field (**Fig. S9**). In the twentieth century, mathematical modeling in oncology was concentrated largely in medical physics, radiobiology and probabilistic models of somatic evolution. Over the past 25 years the field has expanded around spatial, ecological and multiscale modelling approaches that connect tumour progression with the tumour microenvironment. This shift has shaped the contemporary landscape of Mathematical Oncology, consolidating a view of cancer as a spatially structured eco-evolutionary process.

## Discussion

In this work, we provided a comprehensive analysis of mathematical modeling in oncology based on an LLM-curated bibliometric dataset comprising more than 14,000 publications over 140 years. Combining quantitative analysis with LLM-assisted curation (**Fig. 1, 2**), we examine the field’s geographic (**Fig. 3**) and temporal evolution (**Fig. 4**), and characterize its thematic development both globally (**Fig. 5**) and across distinct research lines (**Fig. 6**).

In agreement with the recent scientific investigation^37^ we show that LLMs are capable of curating scientific references at the same level of experienced scientists (**Fig. 1c**). Despite differences between foundational models, the minimal balanced accuracy is 0.75 (GPT-mini, Naive), demonstrating remarkable capabilities even for underperforming models. Overall, Claude displayed the best performance across foundational models (**Tab. S4**), reaching the top position with Claude Sonnet-4.6 (TWiMO-based).

It is well known that there is a trade-off between context and performance of LLMs, where either too little or too much context can undermine performances on an assigned task^39^ Our results confirm the presence of this trade-off for scientific metadata curation, with the TWiMO-based strategy resulting with the best performance in general, but not in all cases Of note, in this study we always employed the same model during the curation and the context generation (see **Methods**), so the results may be affected not only by the presence or absence of the context, but by the quality of the context generated by different models.

Using LLMs to curate verified scientific references, rather than generate *de novo* scientific text, may be essential for their responsible integration into research workflows beyond bibliometric analysis. As scientific output continues to expand, approaches needed to retrieve, summarize, and organize evidence at scale. Yet, LLMs remain vulnerable to hallucinations^38^ including fabricated citations, which are particularly problematic in scientific applications. Here, we demonstrate a simple and flexible LLM-based curation pipeline grounded in real citations and therefore inherently protected from hallucinated references. This workflow can be applied to any scientific dataset, for any domain of science. Here, demonstrate its application to characterize the evolution of mathematical modeling in oncology.

The LLM-curated dataset provides an unprecedented view on the countries involved in mathematical modeling in oncology. Geographic patterns (**Fig. 3 Fig. S2**) confirm the leading role of U.S. and U.K. institutions while revealing increasing contributions from Europe, the Middle East, and Asia, particularly China. The comparison between institutional (**Fig. 3e**) and sponsor (**Fig. 3f**) rankings suggests close link between nstitutional strength and targeted funding. Notably, the U.S. and the U.K. are the only countries reporting to provide clinically oriented funding, which likely contributes to their prominence among leading institutions. This observation, however, should be interpreted with caution, as detailed funding information was available for only 48% of the entries, limiting the completeness of the analysis.

The broader international reach is mirrored by a steady rise in publications on mathematical modeling in oncology (**Fig. 4a**). Earlier reports attributed this growth to advances in Systems Biology and new modeling approaches^11^ The analysis of terminology and key contributors (**Figs. 5–6 S7, S8, S9**) indicates that it also reflects increasing attention to therapy over cancer biology, and increasing relevance of spatial modeling characterizing evolving, living ecosystem.

These trends indicate that multidisciplinarity and a growing focus on clinical applications have been major drivers of the field’s expansion. Journal subject areas corroborate this view: over half of the publications fall within Medicine, Biochemistry/Genetics, and Mathematics (**Fig. 3c**), with a stable distribution over time (**Fig. S3**). An independent bibliometric study on a smaller sample^50^ similarly highlighted the field’s interdisciplinarity, reinforcing its centra role in shaping mathematical modeling in oncology. The journal analysis reveals the strong scientific impact of mathematical modeling in oncology, with 63% of the papers published in Q1 more than double the expected 30%^41^ (**Fig. 3b**). Citation patterns also indicate that publicly available publications (**Fig. S4**) achieve greater impact, with Bronze OA (see **Methods**) showing the highest total citations and Green OA (see **Methods**) the highest annual citation rate (**Table S1**). The strong performance of Green OA underscores the value of preprint-based accessibility in increasing research visibility.

Analysis of the most represented AKs reveals a clear shift in mathematical modeling in oncology from tumor-biology themes toward therapy-focused topics (**Fig. 5b**). Terms related to specific tumors and cancer therapies are strongly represented in the LLM-curated dataset (**Fig. 5a**) and temporal trends show declining use of terms related to tumor biology (**Fig. S5c**). The popular AKs in the author clusters (**Fig. 6 S7**), show a similar trend, with a strong relevance of terms related to therapy (e.g. *Chemotherapy*”, *Radiotherapy*”, *Adaptive Therapy* in **Fig. S7**) and a increase of their use in time across multiple research groups in parallel (**Fig. S9**).

This trend reflects the growing integration of mathematical and quantitative approaches into clinical oncology, which, despite some limitations, are increasingly offering mechanistic insights to improve cancer treatment^34^ This shift appears to focus on breast cancer and tumors of the genitourinary, gynecologic, and central nervous systems, indicating uneven attention across cancer types (**Fig. 5i, j, S6**). While traditional modeling approaches, like differential equations, PK/PD, and probabilistic models, remain dominant in mathematical modeling in oncology, temporal trends reveal growing contributions from other techniques, including ABMs and EGT (**Fig. 5h**). A relevant increase is also observed for publications incorporating ML (**Fig. 5a, d**), possibly driven by the emergence of Mechanistic Learning^29^

Comprehensive bibliographic analyses indicate that the reorientation toward clinical applications is a key factor in the emergence of Mathematical Oncology as a distinct subfield. Thematic clusters (**Fig. 5c**) show that “PK/PD,” “Systems Biology,” and “Mathematical Oncology” occupy separate semantic domains, with the latter focused on therapy, optimization, and data-driven approaches. Temporal trends reveal that “Mathematical Oncology” has shown the most prominent growth among the terms related to scientific (**Fig. S5d**), and the majority of author clusters (**Fig. 6**) employ this term to describe their research. This increased adoption occurs in parallel to the increasing prominence of therapeutic applications and spatial modeling, remarking the importance of mathematical modeling to characterize complex, nonlinear interactions in space and time.

Our temporal analysis confirms that Mathematical Oncology has fully embraced the importance that the microenvironment plays in cancer progression (**Fig. 6, Fig. S9**), providing cancer research with a essential tool to understand tumor evolution and the effect of therapies within the lens of mathematical modeling.

In conclusion, our analysis addresses fundamental gaps in interpretation of mathematical modeling in oncology, offering a comprehensive and quantitative perspective on its evolution over 140 years. Our AI-assisted curation framework achieved high precision and dataset quality, highlighting its potential for scalable reference curation beyond bibliometric analyses. Mathematical Oncology is a thriving interdisciplinary field with a distinct scientific identity, which cannot be merely described as the se of mathematics in cancer research. We propose to define Mathematical Oncology as the use of interpretable mathematical models, integrating clinical, biological and physical knowledge, to enhance cancer screening, understand disease evolution, guide therapy, and strengthen forecasting.

## Materials and Methods

### Datasets

#### This Week in Mathematical Oncology dataset (TWIMO dataset)

The complete list of publications featured in *This Week in Mathematical Oncology* (TWIMO) is maintained and updated weekly by the editors on GitHub (**Table S2**). From this list, we retrieved the digital object identifiers (DOIs) of all entries published until 2024, and extracted the corresponding bibliographic information from Scopus for analysis (**Table S3**).

#### Keyword-based dataset

To ensure inclusiveness in our study, we derived a comprehensive dataset through keyword-based search in Scopus. As with most scientific databases, Scopus enables the use of logical operators to combine search terms. We therefore designed a comprehensive query integrating terminology commonly used in both cancer research (e.g., “Cancer,” “Tumor,” “Neoplasm”) and mathematical modeling (e.g., “Mathematical Model,” “Mathematical Oncology,” “Computational Oncology,” “PK/PD”) (**Table S3**). We selected all citable peer-reviewed document types (articles, reviews, and conference papers) published in complete years up to 2025, since the analysis was conducted in 2026. Additionally, all documents explicitly containing the field-defining terms “Mathematical Oncology”, “Computational Oncology”, and “Cancer Systems Biology” included, while those referring to “Molecular Dynamics” excluded to avoid false positives identified in preliminary iterations of our analysis. This search yielded a dataset comprising 36,010 publications. Manual curation of 200 sample publications from this dataset resulted in 66% of the items deemed relevant for at least 2 over three experts, and 50.5% of the publications considered relevant by all the experts (see **Human Curation**).

#### Data cleaning

The most relevant metadata for analysis, i.e. authors names, AKs, institutions, and journal titles, were harmonized using a field-specific, thesaurus-based cleaning workflow. For each field, we identify possible lexical variants and spelling mistakes, we cluster terms with a similar Levenshtein distance (i.e. terms differing of a few characters). These clusters were used to automatically generate a thesaurus file, which was manually curated to map each possible synonym to unique term. Additional ad-hoc standardization applied to fields with high variability, particularly affiliations (e.g., removal of department-level fragments and consolidation of institutiona name variants) and keywords (e.g., each instance of “Modelling” was replaced with “Modeling”, each instance of “Tumour” with “Tumor”, etc.). When a unique identifier was available (e.g. for authors, which are uniquely identified in Scopus by a specific code), we used it to further minimize duplicates. This hybrid strategy, combining automated detection with curated normalization, reduced terminological redundancy while preserving domain-relevant meaning, thereby improving consistency and robustness for downstream bibliometric analyses.

### Human Curation

Three independent experts curated a sample of 200 randomly selected publications (validation dataset) derived from the keyword-based dataset. This validation dataset was used for estimating the number of false positives in the keyword-based dataset and to validate the performance of the LLM-curation (see **LLM Curation**). Each expert evaluated all the documents, assessing their relevance for the broad field of mathematical oncology. More precisely, each expert was asked to answer true or false to the following question: “Please evaluate this paper according to the following definition of mathematical oncology: the use of interpretable mathematical models—integrating clinical, biological and physical knowledge— to enhance cancer screening, understand disease evolution, guide therapy, and strengthen forecasting”.

Notice that the instructions provided, despite mentioning specifically mathematical oncology, include a definition that does not exclude adjacent fields such as Cancer Systems Biology, or PK/PD modeling when applied to cancer The results of the human curation are provided in **Figure S1**

### LLM Curation

LLM curation was performed on all 36,010 entries of the keyword-based dataset, testing different foundational models (Claude, Gemini 3.1, GPT 5.4) in different model sizes (for Claude: Opus 4.6, Sonnet 4.6, Haiku 4.5; for Gemini 3.1: Pro, Flash, Flash-Lite; for GPT 5.4: regular, mini, and nano). Given the importance of context in LLMs performance^39^ we also tested two different context strategies: Naive and TWiMO-based. For each model and each strategy, the LLM-curator was asked to: 1) provide a score from zero to one indicating how likely the publication could be of interest for a mathematical oncologist, and one sentence motivating its decision (see **Supplementary Appendix 2, Prompt 2.1**)

#### Context Strategies

The Naive strategy provides no context to the LLM Curator outside the Prompt 2. (see **Fig. 2**). The TWiMO-based strategy follows three steps of context generation: 1. Documents Retrieval; 2. Summarization; 3. Synthesis. The objective of this pipeline is to automatically produce instructions for the LLM curator to perform its task.

In the document retrieval, we select all the reviews from the TWiMO dataset and we download their full text using OpenAlex. Fetching with OpenAlex resulted in successful retrieval of 88 reviews, corresponding to 35.3% of all the reviews contained in the TWiMO dataset. In the summarization step, we provide each of the selected documents as input to an LLM summarizer, asking to generate a summary of the documents under 300 words (**Supplementary Appendix 2, Prompt 2.2**). In the synthesis step, we compose the summaries generated by the LLM Summarizer in a single prompt to a LLM Synthetizer, with the task to produce a "synthesis" of all the summaries. We specify to the LLM Synthesizer to provide the synthesis in the form of concise guidelines(<500 words) to follow in order to assess f paper could be of interest for a mathematical oncologist (**Supplementary Appendix 2, Prompt 2.3**). n conclusion, the synthesis is provided as a configuration prompt to the LLM curator to perform its task (**Supplementary Appendix 2, Prompt 2.4**).

Importantly, when using the TWiMO-based strategy we use the same LLM model for all the steps of the curation. For instance, when testing Claude Sonnet 4.6 (best LLM curator), all the summaries and the synthesis are automatically generated using the same exact model. The synthesis and one of the summaries generated by Claude Sonnet 4.6 are provided in **Supplementary Appendix 3, LLM Output 3.1** and **3.2**

#### Performance evaluation

Each model and each strategy was compared with the human annotations of the validation dataset, considering the majority voting (2/3 evaluators voting true) as ground truth. To account for the inherently random nature of LLMs, we executed the LLM curation on the validation dataset five times for each model.

For each model we evaluated the mean and the standard deviation of different metrics: area under the curve of the receiver-operator characteristic (ROC AUC), average precision (Mean PR), F1 score, and balanced accuracy (BA). For each metric, we derived the mean and the standard deviation for the best threshold T to discriminate between positive (LLM score above T) and negative (LLM score below T) documents.

To account for the imbalance of the validation dataset (66% positives), we selected the LLM curator with the highest balanced accuracy (Claude Sonnet 4.6 with the TWiMO-based strategy), which also resulted in the best F1 score Of note, Claude Sonnet 4.6 resulted also in the best ROC AUC and average precision using the Naive strategy, denoting excellent internal representation of the model for concepts related to mathematical modeling in oncology. A complete overview of the performances of every mode, for each context strategy is provided in **Table S4**

### Rankings

All the rankings displayed in the manuscript (e.g. authors, countries) have been programmatically produced using Python and Matplotlib. Whenever possible, unique identifiers used to prevent duplication spelling inconsistencies Specifically, authors, journals, and institutions identifiers were matched via Scopus IDs, which ensured unambiguous aggregation of entities. Since AKs lack unique identifiers, terms were standardized by capitalization, and eventual synonymous terms were merged together (see **Table S5** for the list of synonyms). For example, “Mathematical Model,” “Mathematical Modeling,” and “Mathematical Modelling” were considered under a single keyword entry.

When it comes to author rankings (e.g. **Fig. S8a**) it is important to underscore that our analysis is limited to publications within the LLM-curated dataset. Our aim is to capture specific contributions to mathematical applications in cancer rather than each author’s broader output outside of mathematics and cancer, thus the total number of publications associated with each researcher do not correspond to their total number of publications.

### All Science Journal Classification (ASJC) areas

To measure the multidisciplinarity of the publications in the query-based dataset, we leveraged the distribution of the ASJC areas. In Scopus, each journal is associated with one or more scientific areas following the ASJC. Since each document in the query-based dataset has been published in a journal, this allowed us to map each individual publication to one or more scientific areas, and to observe the overall distribution in the dataset (**Fig. 3c, Fig. S3**).

### Citations, quartile, and open access

Citation data were obtained from Scopus (**Table S3**). To calculate citation rates per year, total citations were divided by the number of years since publication (e.g., a paper with 100 citations published in 2015 has 10 citations per year in 2025). Scopus also reports Open Access (OA) status, classifying publications into five categories: Not OA (N/A), Gold OA (published in an OA-only journal), Hybrid Gold OA (journal allows OA or non-OA publishing), Green OA (journal publication plus deposition in a public repository such as arXiv or bioRxiv), and Bronze OA (journal provides temporary OA). These classifications enabled stratified analyses of citation distributions by OA type (**Fig. S4**). Kolmogorov-Smirnoff test with Bonferroni correction (when comparing more than two groups) was used to establish statistical significance between different OA modalities (**Table S1**).

Journal quartiles derived from Elsevier’s Scival database (**Table S5**). Quartiles defined using the distribution of Scopus CiteScore™, which reflects the average citations per document over the preceding four years. Journals are assigned to subject fields and classified into quartiles accordingly, with Q1 representing the highest-performing journals. Since quartile data are available only from 2014 onward, analyses were limited to publications from 2014–2024.

### AK Categories

For the analysis of the AK categories, we manually divided the top 100 AKs into 5 categories: Therapy, Biology, Modeling, Cancer Type, and Scientific Area (**Fig. S5a**). When unsure, we labeled the AK as “other”; for instance, the AK “Immune System” could be related to either the biology of the immune system, or with immunotherapy. Thus, we left the AK unlabeled.

### Analysis of Scientific Areas

To analyze the terminology used by the three distinct scientific n mathematical modeling in oncology, i.e. Mathematical Oncology, Systems Biology, and PK/PD Modeling, we selected all the documents in the dataset explicitly mentioning one of these fields in their AKs. Given the different possible spellings for PK/PD modeling, we used a group of regular expressions to match the documents mentioning a set of terms specifically associated with this scientific area (e.g. *“Physiologically Based Pharmacokinetic”* “Pharmacokinetics & Pharmacodynamics”, etc.). Our approach allowed us to identify the documents explicitly positioned by the authors as part of one or (more rarely more) scientific areas However, t presents the limitation of AK-based analysis: not all journals share their AK with scopus leaving about 30% of the documents without any AK. Thus, a minority of the documents explicitly associated to one scientific area could have been missed by our analysis.

### Text mining and regular expression analysis

Regular expressions (regexes) were designed to identify key modeling approaches and cancer types within publication titles, abstracts, and keywords. Regexes representing modeling methods (45 total, **Table S6**) were grouped into seven categories: Differential Equations (22 terms), Agent-Based Models (3), Probabilistic Models (9), Game Theory (10), Machine Learning (3), Data-Driven Models (3), and PK/PD (3).

To capture cancer-specific research topics, we compiled a list of 197 regexes based on cancer types listed in *Wikipedia* (**Table S7**), organized into 13 categories: Bone and Muscle Sarcomas (10), Brain and Nervous System (26), Breast (13), Endocrine System (15), Eye (5), Gastrointestinal (16), Genitourinary and Gynecologic (30), Head and Neck (9), Hematopoietic (41), Skin (9), Soft Tissue Sarcomas (6), Thoracic and Respiratory (15), and HIV/AIDS-related (2).

For each regex, all titles, abstracts, and keywords were programmatically scanned. A publication was counted once per regex if at least one match was found, regardless of multiple occurrences within the same record.

### Bibliometric Networks

Two types of bibliometric networks were analyzed: (1) a co-occurrence network of AKs (**Fig. 4c**) and (2) bibliographic coupling networks of authors (**Figs. 6, S9**). Both were generated using VOSviewer^35^ and visualized as chord diagrams via the **D3.js** JavaScript library.

In the co-occurrence network, link width represents the frequency with which two AKs appear together in the same publication, and node size corresponds to the total occurrences of each AK. In the bibliographic coupling network, link strength reflects the extent to which two authors cite common documents, thus highlighting shared research trajectories or collaborations ^35^ Node size is proportional to an author’s overall connectivity within the network.

Both networks were clustered using VOSviewer’s internal similarity-based algorithm (see **Appendix S1**). The clustering granularity is controlled by the “resolution” parameter, where higher values yield finer partitions. For the AK co-occurrence network (**Fig. 5c**), a resolution of **0.7** was used; for author coupling networks, **1.1**

## Acknowledgments

**FP JW, and ARAA** gratefully acknowledge funding by the NCI via the Cancer Systems Biology Consortium (CSBC) U54CA274507 and support from the Moffitt Cancer Center & Research Institute’s Center of Excellence for Evolutionary Therapy. **SH** acknowledges Wenner-Gren Stiftelserna/the Wenner-Gren Foundations (WGF2022-0044) and the Kjell och Märta Beijer Foundation. **DAH** acknowledges funding from the Cancer Prevention Research Institute of Texas (RP220225) and the National Science Foundation (DMS 2436499). **GL** acknowledges grant PID2023-146347OA-I00 funded by MICIU/AEI/10.13039/501100011033 and ERDF/EU, well as grant RYC2022-036010-I funded by MICIU/AEI/10.13039/501100011033 and ESF+. **MS** acknowledges Schmidt Sciences, LLC. **KG** gratefully acknowledges funding by BMFTR as “SATURN3: Spatial And Temporal Resolution of Intratumoural Heterogeneity in 3 hard-to-treat CaNcers”, Project number 01KD2206L. and the Bruno and Helene Jöster Foundation as “CLONETRAC Tracking the clonal dynamics of cancer through treatment at the single-cell level. We used generative artificial intelligence (Grammarly, ChatGPT 5.2) to proofread some sections of the manuscript, but never to generate new text or images and the authors take responsibility for the final content within the manuscript.

## Data Availability Statement

The data generated in this study are publicly available on Zenodo (DOI: https://doi.org/10.5281/zenodo.20545347).

## Author contributions (CRediT)

Conceptualization, Investigation, Formal Analysis, Methodology, Project Administration, Writing Original Draft Preparation: **FP & JW** Data Curation: **FP, MS, GL, JW** Funding Acquisition: **JW & ARAA** Software: **FP** Supervision: **JW** Visualization: **FP, JW, MS, GL** Writing Review & Editing: **MS, SM, FdK, AB, KG, GL, DAH, SH, DB, ARAA.**

