## Supplementary Material for "140 Years of mathematical modeling in oncology through AI-assisted curation"

Franco Pradelli[1*], Maximilian Strobl[2,3,4,5], Sadegh Marzban[1], François de Kermenguy[6], Ari Barnett[1], Katyayni Ganesan[7], David A. Hormuth II[8], Sara Hamis[9], Dhananjay Bhaskar[10], Guillermo Lorenzo[8,11], Alexander R. A. Anderson[1], Jeffrey West[1*]

1. Integrated Mathematical Oncology Department, Moffitt Cancer Center, Tampa, FL. USA.
2. Department of Genomic Medicine, Cleveland Clinic Research, Cleveland Clinic, Cleveland, USA.
3. Department of Life Sciences, Imperial College, London, UK.
4. Institute of Cancer Research, London, UK.
5. I-X Centre for AI in Science, Imperial College London, London, UK.
6. Harvard Medical School, Dana-Farber Cancer Institute, Brigham and Women's Hospital, Boston, MA. USA.
7. Institute of Computational Cancer Biology (ICCB), University Hospital and University of Cologne, Cologne, Germany.
8. Oden Institute for Computational Engineering and Sciences, The University of Texas at Austin, Austin, TX, USA.
9. Division of Systems and Control, Department of Information Technology, Uppsala University, Uppsala, Sweden.
10. Biomedical Engineering Department, University of Wisconsin-Madison, Madison, WI, USA.
11. Group of Numerical Methods in Engineering, Department of Mathematics, and CITEEC, University of A Coruña, A Coruña, Spain.

*Corresponding Authors

**This file contains:**

- Supplementary Appendix 1: Clustering in Bibliographic Networks using VOSViewer
- Supplementary Appendix 2: Prompts
- Supplementary Appendix 3: LLM Output
- Supplementary Tables from S1 to S8
- Supplementary Figures from S1 to S9

### Appendix 1: Clustering in Bibliographic Networks using VOSViewer

The following is meant to provide a brief explanation of the clustering algorithm in VOSViewer to understand how and when two nodes in a network are considered “similar”. For additional information, see[1].
As an example of the clustering technique, consider the case of a co-occurrence network (as those we generated for the author keywords in our datasets). Given a set of terms T of n elements and a set of documents D of m elements, we can define the co occurrence matrix C of the terms in A as the square symmetric matrix where each element is:

$c_{ij}$ = Number of documents where the term i and the term j occur together

Notice that each term corresponds to a row (or a column) of the matrix C, which is a vector of the co-occurrences with all the other terms. Also notice that $c_{ii}$ correspond to the number of total occurrences of the term across all documents.
Now, how can we define the similarity between two terms using the information of their co-occurrence profile? This is the core question dissected in the publication[2], and there are actually different methods to measure such similarity. The method used in VOSViewer uses the following similarity metric:

$$s_{ij} = \frac{mc_{ij}}{c_{ii}c_{jj}}$$

Where m stands for the total sum of the occurrences of all terms (i.e. $m = \sum_{k=1}^{n} c_{kk}$). The reason why this metric works can be easily understood following a probabilistic approach. Given a term i and a term j, the probability of each of these terms to appear in a given document is:

$$p_i = \frac{c_{ii}}{m} \qquad p_j = \frac{c_{jj}}{m}$$

It follows that the probability to observe them in the same document by simple chance is

$$p_{ij} = p_i p_j = \frac{c_{ii}c_{jj}}{m^2}$$

While the actual frequency where the two terms are observed together in the documents is:

$$f_{ij} = \frac{c_{ij}}{m}$$

So, the similarity metric used in VOSViewer can be thought as a ratio between the actual frequency of co-occurrence between two terms and the probability they would occur together by chance:

$$s_{ij} = \frac{f_{ij}}{p_{ij}} = \frac{mc_{ij}}{c_{ii}c_{jj}}$$

So, if two terms have a high frequency of co-occurrence despite a low probability of co-occurring together, their similarity index will be high. If two terms have a low frequency of co-occurrence and a high probability of occurring together (e.g. because they are both common terms), their similarity will be low.
The clusterization algorithm employed by VOSViewer simply tends to group terms based on their, maximizing the following function[1]:

$$V = \sum_{i<j} \delta(g_i, g_j)(s_{ij} - \gamma)$$

Where $g_i$ and $g_j$ are two clusters, $\boldsymbol{\delta}$ is a function which is 1 when $g_i$=$g_j$ and 0 otherwise, and $\boldsymbol{\gamma}$ is a resolution parameter which regulates the granularity of the clustering. The higher is $\boldsymbol{\gamma}$, the higher will be the number of clusters. In all the clusters we generated, we used the default VOSViewer value for $\boldsymbol{\gamma}$, which is 1.

### Appendix 2: Prompts

This section lists all the prompts adopted for every application of LLMs in the manuscript.

I am providing you the content of a json file containing a list of publications with the following metadata: Title, Author full names, Source title, Author Keywords, DOI, Year, Abstract, ID

JSON:
<JSON BATCH>

Give a score from 0 to 1 indicating how likely each publication could be of interest for Mathematical Oncologists.
Also provide a concise motivation (one sentence) on the reasoning.
Your answer should be in the following json format, where ID is the ID of each publication, INTEREST_SCORE the score you provided, and MOTIVATION the motivation you provided:
{ID: {\"interest_score\": INTEREST_SCORE, \"motivation\": \"MOTIVATION\"}, ...}.

**Prompt 2.1. Prompt for LLM curators.** The <JSON BATCH> is replaced at each call of the LLM curator with a json containing the metadata of 10 publications, and the scoring is performed at the same time for all ten. Thus, the keyword-based dataset is divided into 3601 batches in total, and each batch is evaluated independently. This prompt is provided as input to all LLM-curators, regardless the context strategy adopted (Naive or TWiMO-based).

Answer like an experienced scientist in the field of mathematical modelling in cancer research.

Provide a concise summary of the main findings of the review paper below, focusing on the use of mathematical models in cancer research. The sumary should be in bullet points and should not exceed 300 words. Here is the content of the paper (parsed with pyPDF):

<PDF TEXT>

**Prompt 2.2. Prompt for LLM summarizer.** The <PDF TEXT> is replaced at each call with the text of the review to summarize.

You are provided with a list of summaries of review papers related to the field of Mathematical Oncology.

Your task is to synthesize the key themes and findings across all the review summaries in the form of recommendations to follow in order to assess if a paper could be of interest for a fellow mathematical oncologist. The synthesis should be no longer than 500 words and include positive and negative indicators of interest. Here are the review summaries:

<SUMMARIES_LIST>

**Prompt 2.3. Prompt for LLM synthesiser.** The <SUMMARIES LIST> is replaced at each call with the list of the summaries generated by the LLM summarizer (see **Methods** and **Prompt 2.2**)

You are an experienced scientist in the field of mathematical modelling in cancer research with the task of evaluating scientific publications. In your evaluations, you should rely on the following guidelines:

<guidelines>{ SYNTHESIS }</guidelines>

**Prompt 2.4. Configuration prompt for LLM Evaluator in the TWiMO-based strategy.** The <SYNTHESIS> is replaced at each call with the list of the synthesis generated by the LLM synthesizer (see **Methods** and **Prompt 2.3**)

### Appendix 3: LLM Output

**# Synthesis of Key Themes and Recommendations for Mathematical Oncologists**

**## Core Positive Indicators of Interest**

****Strong relevance signals:****

- ****Explicit mechanistic mathematical frameworks**:** Prioritise papers presenting or critically evaluating ODE/PDE systems, agent-based models (ABMs), evolutionary game theory, stochastic models, or QSP platforms applied to tumour or immune dynamics. Reviews from this corpus covering adaptive therapy, eco-evolutionary modelling, and tumour-immune interactions (e.g., papers on evolutionary game theory, QSP for immuno-oncology, leukaemia modelling, prostate cancer) exemplify high-value targets.

- ****Quantitative biological predictions with clinical translation****: Papers linking mathematical model outputs directly to patient outcomes — such as PSA-guided adaptive therapy models achieving 81% clinical prediction accuracy, or TGI-OS joint models informing dose selection — indicate directly actionable modelling work.

- ****Multi-scale or hybrid modelling frameworks****: Reviews integrating intracellular signalling, cellular dynamics, and tissue/organ-scale behaviour (e.g., PhysiCell-based platforms, mechanistic learning combining ML with mechanistic models, digital twins) represent methodologically sophisticated and translatable contributions.

- ****Eco-evolutionary dynamics and resistance modelling****: Papers explicitly deploying Lotka-Volterra, replicator dynamics, Stackelberg games, or phenotype-structured PDEs to analyse competitive suppression, adaptive therapy, or collateral sensitivity are of high intrinsic relevance.

- **Novel mathematical formalisms applied to cancer**: Information-theoretic frameworks (Shannon entropy, mutual information), topological data analysis (persistent homology), and Waddington landscape formalisms applied quantitatively to tumour biology represent emerging methodological contributions warranting attention.

- **Parameterisation from imaging or molecular data**: Reviews describing model calibration using quantitative MRI, PET, ctDNA dynamics, or single-cell omics — particularly where patient-specific predictions are validated — indicate strong translational readiness.

**## Negative Indicators (Deprioritise)**

- **Purely biological or clinical narrative reviews**: Papers without mathematical content, quantitative predictions, or explicit modelling frameworks (e.g., purely mechanistic descriptions of the Warburg effect, immunoediting biology, or liquid biopsy clinical workflows) offer limited direct value unless serving as parameterisation sources.

- **Methodological papers tangential to oncology**: Reviews on covariate modelling best practices, computational biology career development, or biologist-computer scientist collaboration guidelines have negligible mathematical oncology content.

- **Incorrect document parsing**: Several summaries indicate PDF parsing errors returning unrelated documents; these should be immediately excluded.

- **Descriptive clinical epidemiology**: Systematic reviews of topographic brain metastasis distributions or purely observational cancer biology, absent spatial modelling or quantitative frameworks, warrant low priority.

- **Broad computational biology overviews**: Papers addressing general data science infrastructure, FAIR principles, or omics data management without cancer-specific mathematical modelling content are peripheral.

#### Overarching Recommendations

A paper merits close reading if it: (1) proposes, validates, or critiques a mathematical/computational model with biological mechanistic grounding; (2) connects model outputs to experimentally or clinically measurable quantities; (3) addresses treatment optimisation, resistance evolution, or tumour-immune dynamics quantitatively; or (4) introduces methodological innovations (novel model classes, hybrid approaches, uncertainty quantification) applicable across cancer types. Papers scoring on multiple criteria simultaneously — mechanistic rigour, clinical data integration, and therapeutic implication — represent the highest priority reads for a practising mathematical oncologist.

**LLM Output 3.1. Synthesis (or guidelines) produced by Claude Sonnet 4.6 for the TWiMO based strategy**. The output is in Markdown (.md) format

Title: Digital twins for health: a scoping review
Authors: Katsoulakis, Evangelia (35763630300); Wang, Qi (57210440107); Wu, Huanmei (18438486400); Shahriyari, Leili (56179571000); Fletcher, Richard (7201875267); Liu, Jinwei (56487079400); Achenie, Luke (7003325038); Liu, Hongfang (7409753328); Jackson, Pamela (56266947800); Xiao, Ying (58951602200); Syeda-Mahmood, Tanveer (7003810087); Tuli, Richard (7003452100); Deng, Jun (36820468900)
Journal: npj Digital Medicine
Year: 2024
DOI: 10.1038/s41746-024-01073-0

Summary:
**## Summary: Digital Twins for Health – Key Findings Relevant to Mathematical Modelling in Cancer Research**

**### Definition & Framework**
- A Digital Twin for Health (DT4H) is defined as a virtual representation of a patient enabling dynamic simulation of treatment strategies, health trajectory prediction, and early intervention, integrating multi-scale, multi-modal data (clinical, genomic, molecular, environmental)

**### Cancer-Specific Mathematical Modelling Applications**
- ****Oropharyngeal cancer****: Patient-physician digital twin dyads using deep Q-learning successfully predicted optimal sequential vs. concurrent chemoradiation treatment selection
- ****Triple-negative breast cancer (TNBC)****: Quantitative MRI combined with biologically-based mathematical modelling accurately predicted neoadjuvant chemotherapy responses
- ****Non-small cell lung cancer (NSCLC)****: Virtual clinical trials simulating >25,000 lesion measurements from >500 patients informed optimal salvage therapy decisions following pembrolizumab progression
- ****Metastatic spine cancer****: DT-based finite element modelling predicted vertebral fracture risk following stereotactic body radiotherapy (SBRT)
- ****Childhood malignancies (PRIMAGE)****: Multiscale in silico analytics integrating imaging biomarkers support personalised diagnosis and prognosis

**### In Silico Clinical Trials**
- Mathematical models enable synthetic control arms, reducing costly patient recruitment; validated in breast cancer (palbociclib), lung cancer (alectinib), and leukemia (blinatumomab)

- VICTRE trial demonstrated strong concordance between in silico and traditional clinical trial outcomes for breast lesion detection

**### Key Challenges**
- Integration of multi-scale biological data into coherent mathematical models remains technically demanding
- Data quality, bias, and privacy concerns limit model accuracy and generalisability
- Lack of standardised interoperability frameworks across institutions

**### Outlook**
- Convergence of AI/ML, mechanistic modelling, and real-time patient data holds strong promise for predictive, personalised oncology DTs

**LLM Output 3.2. Example of one review summary produced by Claude Sonnet 4.6 for the TWiMO based strategy**. The output is in Markdown (.md) format.

**Example input (JSON):**

{
"Title": "Modelling physiological and pharmacological control on cell proliferation to optimise cancer treatments",
"Authors": "Clairambault J.",
"Author full names": "Clairambault, J. (6602523420)",
"Author(s) ID": "6602523420",
"Year": 2009,
…
}

**Example output (JSON):**

{
"interest_score": 0.75,
"motivation": "Comprehensive review of physiologically structured PDE models for cell proliferation integrating PK/PD, circadian control, and optimal drug delivery, directly relevant to mathematical oncology and treatment optimisation."
}

**LLM Output 3.3. Example of one evaluation produced by Claude Sonnet 4.6 with the TWiMO based strategy**. To simplify the representation, we show a single publication as input and a single output. To optimize token utilization and response timing, our architecture divides the publications into batches of 10 publications that are evaluated at the same time (see **Methods**).

### Supplementary Tables

| Test | Median Citations | p value | α value | Significant | Median Citations per Year | p value | α value | Significant |
|---|---|---|---|---|---|---|---|---|
| OA vs NOA | OA: 12.0<br>NOA: 11.0 | $2.44×10^{−12}$ | 0.05 | True | OA: 1.9<br>NOA: 1.00 | $4.39x10^{-122}$ | 0.05 | True |
| Gold OA vs Green OA | Gold OA: 10.0<br>Green OA: 12.0 | $4.99×10^{−7}$ | 0.0083 | True | Gold OA: 1.90<br>Green OA: 2.00 | $4.79x10^{-4}$ | 0.0083 | True |
| Gold OA vs Bronze OA | Gold OA: 10.0<br>Bronze OA: 25.0 | $1.47×10^{−57}$ | 0.0083 | True | Gold OA: 1.90<br>Bronze OA: 2.2 | $1.14x10^{-7}$ | 0.0083 | True |
| Gold OA vs Hybrid Gold OA | Gold OA: 10.0<br>Hybrid Gold OA: 10.0 | 0.48 | 0.0083 | False | Gold OA: 1.9<br>Hybrid Gold OA: 1.8 | 0.62 | 0.0083 | False |
| Green OA vs Bronze OA | Green OA: 12.0<br>Bronze OA: 25.0 | $7.58×10^{-39}$ | 0.0083 | True | Green OA: 2.0<br>Bronze OA: 2.2 | $8.58\ x10^{-4}$ | 0.0083 | True |
| Green OA vs Hybrid Gold OA | Green OA: 12.0<br>Hybrid Gold OA: 10.0 | $8.84×10^{−5}$ | 0.0083 | True | Green OA: 2.0<br>Hybrid Gold OA: 1.8 | $5.42x10^{-4}$ | 0.0083 | True |
| Bronze OA vs Hybrid Gold OA | Bronze OA: 25.0,<br>Hybrid Gold OA: 10.0 | $8.84×10^{-42}$ | 0.0083 | True | Bronze OA: 2.2<br>Hybrid Gold OA: 1.8 | $5.89x10^{-6}$ | 0.0083 | True |

**Table S1. Results of the Kolmogorov–Smirnov tests comparing citation distributions across open-access categories.**The first column (“Test”) specifies the pairwise comparison of open-access modalities. The left panel (grey) summarizes analyses based on total citations per document and reports, for each comparison, the median citation count of the two categories, the Kolmogorov–Smirnov p-value, the significance threshold (α), and the resulting significance decision. The right panel (white) presents analogous results for citations per year. A default significance threshold of α = 0.05 is applied; when a different α value is reported, it reflects the use of a Bonferroni correction for multiple testing.

| Resource | Link |
|---|---|
| The Mathematical Oncology Website | https://mathematical-oncology.org/ |
| The Mathematical Oncology Newsletter (“This Week in Mathematical Oncology”) | https://thisweekmathonco.substack.com/ |
| Collection of Papers in “This Week in Mathematical Oncology” | https://github.com/MathOnco/Newsletter-Bibliography |

**Table S2. Online resources related to TWIMO.** The references we used for our TWIMO dataset can be found in the “Collection of Papers in “This Week in Mathematical Oncology".

1

| | |
|---|---|
| **Query** | (<br>PUBYEAR < 2026<br>)<br>AND<br>(<br>DOCTYPE(ar)<br>OR DOCTYPE(cp)<br>OR DOCTYPE(re)<br>)<br>AND<br>(<br>LANGUAGE("english")<br>)<br>AND<br>(<br>(<br>"Mathematical Oncology"<br>OR "Cancer Systems Biology"<br>OR "Computational Oncology"<br>)<br>OR<br>(<br>TITLE-ABS-KEY<br>(<br>"Cancer*"<br>OR "Tumor*"<br>OR "Tumour*"<br>OR "Neoplas*"<br>OR "Carcino*"<br>)<br>AND<br>TITLE-ABS-KEY-AUTH<br>(<br>"Mathematical model*"<br>OR "Agent based model*"<br>OR "Stochastic model*"<br>OR "Numerical model*"<br>OR "Deterministic model*"<br>OR "Game theor*"<br>OR "differential equation*"<br>OR "in silico model*"<br>OR "Cellular Automa*"<br>OR "Computational Oncolog*"<br>OR "Biomechanical Model*"<br>OR "Biomechanistic Model*"<br>OR "Biology based Model*"<br>OR "Mechanism based model*"<br>OR "Biology based Mathematical Model*"<br>OR "PK/PD"<br>OR "PK-PD"<br>OR "Pharmacokinetic/Pharmacodynamic"<br>OR "Pharmacokinetic-Pharmacodynamic"<br>OR "Physiologically based pharmacokinetic model*"<br>OR "Moran proces*"<br>OR "Moran model*"<br>OR ("growth" AND "law")<br>)<br>)<br>)<br>AND NOT<br>(<br>"Molecular Dynamics"<br>) |
| **Dataset size** | 36010 |

| |
|---|
| **Table S3. Query used for deriving the keyword-based dataset.** The search is not case-sensitive and the character "*" represent every possible group of characters (e.g. "Moran Proces*" matches both "Moran Process" and "Moran Processes"). |

1

| Model | Context | ROC AUC | Average PR | F1 Score | BA | Best Threshold (Balanced Accuracy) |
|---|---|---|---|---|---|---|
| claude-sonnet-4-6 | TWiMO | 0.94537656 | 0.96682661 | 0.9150299 | 0.88146168 | 0.25 |
| gemini-3-flash-preview | Naive | 0.94615642 | 0.96657408 | 0.91006089 | 0.86720143 | 0.78 |
| claude-sonnet-4-6 | Naive | 0.94700312 | 0.96892556 | 0.9079732 | 0.87513369 | 0.44 |
| gemini-3.1-pro-preview | Naive | 0.944541 | 0.96561539 | 0.9076036 | 0.86724599 | 0.8 |
| claude-opus-4-6 | TWiMO | 0.9368984 | 0.96164014 | 0.906636 | 0.86733512 | 0.25 |
| gpt-5.4 | TWiMO | 0.93453654 | 0.96572011 | 0.89897724 | 0.87762923 | 0.408 |
| gpt-5.4 | Naive | 0.92103387 | 0.95516894 | 0.89867727 | 0.85534759 | 0.594 |
| claude-opus-4-6 | Naive | 0.93054813 | 0.96003912 | 0.89819736 | 0.86047237 | 0.386 |
| claude-haiku-4-5 | Naive | 0.91579768 | 0.94744546 | 0.89668703 | 0.84393939 | 0.6 |
| claude-haiku-4-5 | TWiMO | 0.90474599 | 0.93362808 | 0.89434743 | 0.84576649 | 0.31 |
| gemini-3.1-flash-lite-preview | Naive | 0.90981506 | 0.93675008 | 0.88961668 | 0.82887701 | 0.81 |
| gpt-5.4-nano | Naive | 0.89170009 | 0.93229507 | 0.8816641 | 0.82036542 | 0.368 |
| gemini-3-flash-preview | TWiMO | 0.90941399 | 0.93678974 | 0.88159513 | 0.83609626 | 0.39 |
| gpt-5.4-mini | TWiMO | 0.86290589 | 0.90214414 | 0.87553029 | 0.80976064 | 0.628 |
| gemini-3.1-pro-preview | TWiMO | 0.90748663 | 0.93834395 | 0.87531747 | 0.83092692 | 0.2 |
| gpt-5.4-nano | TWiMO | 0.86342469 | 0.90904027 | 0.87127992 | 0.79777184 | 0.216 |
| gemini-3.1-flash-lite-preview | TWiMO | 0.8386475 | 0.88907057 | 0.84681871 | 0.76599822 | 0.49 |
| gpt-5.4-mini | Naive | 0.80954717 | 0.88109911 | 0.84240942 | 0.75385239 | 0.78 |

**Table S4. LLM scorers' performance.** Given the imbalance of our validation set, we selected the model to employ based on Balanced Accuracy. Notably, the model achieving the best performances is Claude Sonnet, regardless of the context. For each metric, we provide the mean across 5 LLM curations. The best threshold is provided only for the BA, our metric of choice for selecting the LLM curator (see **Methods**).

2

| Term | Replaced by |
|---|---|
| Mathematical Modelling | Mathematical Model |
| Mathematical Modeling | Mathematical Model |
| Mathematical Models | Mathematical Model |
| Tumor | Cancer |
| Tumour | Cancer |
| ODE | Ordinary Differential Equation |
| PDE | Partial Differential Equation |
| Ordinary Differential Equations | Ordinary Differential Equation |
| Partial Differential Equations | Partial Differential Equation |
| MRI | Magnetic Resonance Imaging |

| | |
|---|---|
| Cancer Chemotherapy | Chemotherapy |
| Singular Controls | Singular Control |
| Cancer Stem Cells | Cancer Stem Cell |

**Table S5. Synonyms table used for the analysis.** When found as AK, the terms on the left column are replaced by terms on the right column.

| API name | Link | Used for |
|---|---|---|
| Scopus Serial Title | https://api.elsevier.com/content/serial/title/issn/ | Get journals metadata |
| Scopus Abstract | https://api.elsevier.com/content/abstract/doi/ | Access publications metadata |
| Scopus Affiliation Retrieval | https://api.elsevier.com/content/affiliation/affiliation_id/ | Access affiliations metadata |
| Scival | https://api.elsevier.com/analytics/scival/scopusSource/metrics | Access journals metrics (e.g. quartile) |

**Table S7. APIs used for the analysis, and their relative use.**

| General Category | Regex Expression |
|---|---|
| Agent-Based Models | "Agent.Based"<br>"Cellular.Automata"<br>"Individual.Based" |
| Data Driven | "Data.Driven"<br>"Mechanistic.Learning"<br>"Sindy" |
| Differential Equations | "Euler Method"<br>"Runge.Kutta"<br>"Finite Difference"<br>"Finite Element"<br>"Finite Volume"<br>"Differential Equation"<br>"Partial Differential Equation"<br>"Ordinary Differential Equation"<br>" PDE "<br>" ODE "<br>"Stochastic Differential Equation"<br>"Reaction.Diffusion"<br>"Lotka.Volterra"<br>"Predator.Prey"<br>"Phase.Field"<br>"Compartmental"<br>"Gompertz"<br>"Logistic"<br>"Nonlocal Partial Differential Equation"<br>"Integro.Differential Equation"<br>"Fractional Differential Equation"<br>"Fractal" |
| Game Theory | "Game.Theor"<br>"Nash Equilibrium"<br>"Pareto Optimal"<br>"Prisoner's Dilemma"<br>"Chicken.Game"<br>"Stag.Hunt"<br>"Lotka.Volterra"<br>"Predator.Prey"<br>"Public.Goods Game"<br>"Public.Good Game" |

| | |
|---|---|
| Machine Learning | "Machine.Learning"<br>"Artificial Intelligence"<br>"Neural Network" |
| PK/PD | "Pharmacokinetic"<br>"Pharmacodynamic"<br>" PK.PD " |
| Probabilistic Models | "Stochastic.Model"<br>"Monte.Carlo"<br>"Gillespie"<br>"Markov.Chain"<br>"Hidden.Markov"<br>"Bayesian"<br>"Moran.Process"<br>"Wright.Fisher"<br>"Branching.Process" |

**Table S6. Regular Expressions for Mathematical Modeling Approaches**

| General Category | Regex Expressions |
|---|---|
| Bone and muscle sarcoma | "Adamantinoma",<br>"Chondrosarcoma",<br>"Chordoma",<br>"Ewing's sarcoma",<br>"Fibrocartilaginous mesenchymoma of bone",<br>"Leiomyosarcoma",<br>"Malignant fibrous histiocytoma of bone/osteosarcoma",<br>"Myxosarcoma",<br>"Osteosarcoma",<br>"Rhabdomyosarcoma" |
| Brain and nervous system | "Astrocytoma",<br>"Brainstem glioma",<br>"Choroid plexus carcinoma",<br>"Cerebellar astrocytoma",<br>"Cerebral astrocytoma",<br>"Craniopharyngioma",<br>"Ependymoma",<br>"Ganglioneuroma",<br>"Glioblastoma",<br>"Glioma",<br>"Hemangioblastoma",<br>"Medulloblastoma",<br>"Meningioma",<br>"Neuroblastoma",<br>"Neurofibroma",<br>"Oligodendroglioma",<br>"Paraganglioma",<br>"Pineal astrocytoma",<br>"Pineocytoma",<br>"Pineoblastoma",<br>"Pituitary adenoma",<br>"Pilocytic astrocytoma",<br>"Primary central nervous system lymphoma",<br>"Primitive neuroectodermal tumor",<br>"Schwannoma",<br>"Visual pathway and hypothalamic glioma" |
| Breast | "Breast cancer",<br>"Ductal carcinoma in situ",<br>"Inflammatory breast cancer",<br>"Invasive ductal carcinoma",<br>"Invasive lobular carcinoma",<br>"Tubular carcinoma",<br>"Invasive cribriform carcinoma of the breast",<br>"Medullary carcinoma",<br>"Male breast cancer",<br>"Phyllodes tumor",<br>"Mammary secretory carcinoma",<br>"Mucinous carcinoma of the breast",<br>"Papillary carcinomas of the breast" |

| | |
|---|---|
| Endocrine system | "Adrenocortical adenoma",<br>"Adrenocortical carcinoma",<br>"Carcinoid",<br>"Gastrinoma",<br>"Glucagonoma",<br>"Insulioma",<br>"Islet cell carcinoma",<br>"Merkel cell carcinoma",<br>"Multiple endocrine neoplasia syndrome",<br>"Pancreatic cancer",<br>"Parathyroid cancer",<br>"Pheochromocytoma",<br>"Somatostatinoma",<br>"Thyroid cancer",<br>"VIPoma" |
| Eye | "Conjunctival melanoma",<br>"Optic nerve glioma",<br>"Orbital lymphoma",<br>"Retinoblastoma",<br>"Uveal melanoma" |
| Gastrointestinal | "Anal cancer",<br>"Appendix cancer",<br>"Cholangiocarcinoma",<br>"Carcinoid tumor, gastrointestinal",<br>"Colon cancer",<br>"Duodenal cancer",<br>"Extrahepatic bile duct cancer",<br>"Gallbladder cancer",<br>"Gastric (stomach) cancer",<br>"Gastrointestinal carcinoid tumor",<br>"Gastrointestinal stromal tumor",<br>"Hepatoblastoma",<br>"Hepatocellular cancer",<br>"Pancreatic cancer, islet cell",<br>"Rectal cancer",<br>"Small intestine cancer" |
| Genitourinary and gynecologic | "Bladder cancer",<br>"Cervical cancer",<br>"Choriocarcinoma",<br>"Embryonal carcinoma",<br>"Endometrial cancer",<br>"Endodermal sinus tumor",<br>"Extragonadal germ cell tumor",<br>"Fallopian tube cancer",<br>"Gestational trophoblastic tumor",<br>"Kidney cancer",<br>"Leydig cell tumour",<br>"Ovarian cancer",<br>"Ovarian epithelial cancer",<br>"Ovarian germ cell tumor",<br>"Penile cancer",<br>"Prostate cancer",<br>"Renal cell carcinoma",<br>"Renal pelvis and ureter, transitional cell cancer",<br>"Seminoma",<br>"Serous tumour",<br>"Sertoli cell tumour",<br>"Teratoma",<br>"Testicular cancer",<br>"Transitional cell cancer",<br>"Ureter and renal pelvis",<br>"Urethral cancer",<br>"Uterine sarcoma",<br>"Vaginal cancer",<br>"Vulvar cancer",<br>"Wilms tumor" |
| Head and neck | "Esophageal cancer",<br>"Head and neck cancer",<br>"Nasopharyngeal carcinoma",<br>"Oral cancer",<br>"Oropharyngeal cancer",<br>"Paranasal sinus and nasal cavity cancer",<br>"Pharyngeal cancer", |

| | |
|---|---|
| | "Salivary gland cancer",<br>"Hypopharyngeal cancer" |
| Hematopoietic | "Acute biphenotypic leukemia",<br>"Acute eosinophilic leukemia",<br>"Acute lymphoblastic leukemia",<br>"Acute myeloid leukemia",<br>"Acute myeloid dendritic cell leukemia",<br>"AIDS-related lymphoma",<br>"Anaplastic large cell lymphoma",<br>"Angioimmunoblastic T-cell lymphoma",<br>"B-cell prolymphocytic leukemia",<br>"Burkitt's lymphoma",<br>"Chronic lymphocytic leukemia",<br>"Chronic myelogenous leukemia",<br>"Cutaneous T-cell lymphoma",<br>"Diffuse large B-cell lymphoma",<br>"Follicular lymphoma",<br>"Hairy cell leukemia",<br>"Hepatosplenic T-cell lymphoma",<br>"Hodgkin's lymphoma",<br>"Intravascular large B-cell lymphoma",<br>"Large granular lymphocytic leukemia",<br>"Lymphoplasmacytic lymphoma",<br>"Lymphomatoid granulomatosis",<br>"Mantle cell lymphoma",<br>"Marginal zone B-cell lymphoma",<br>"Mast cell leukemia",<br>"Mediastinal large B cell lymphoma",<br>"Multiple myeloma/plasma cell neoplasm",<br>"Myelodysplastic syndromes",<br>"Mucosa-associated lymphoid tissue lymphoma",<br>"Mycosis fungoides",<br>"Nodal marginal zone B cell lymphoma",<br>"Non-Hodgkin lymphoma",<br>"Precursor B lymphoblastic leukemia",<br>"Primary central nervous system lymphoma",<br>"Primary cutaneous follicular lymphoma",<br>"Primary cutaneous immunocytoma",<br>"Primary effusion lymphoma",<br>"Plasmablastic lymphoma",<br>"Sézary syndrome",<br>"Splenic marginal zone lymphoma",<br>"T-cell prolymphocytic leukemia" |
| Skin | "Basal cell carcinoma",<br>"Squamous cell carcinoma",<br>"Squamous cell skin cancer",<br>"Skin adnexal tumors",<br>"Melanoma",<br>"Merkel cell carcinoma",<br>"Keratoacanthoma",<br>"Sarcomas of primary cutaneous origin",<br>"Lymphomas of primary cutaneous origin" |
| Soft Tissue Sarcoma | "Angiosarcoma",<br>"Fibrosarcoma",<br>"Liposarcoma",<br>"Malignant peripheral nerve sheath tumor",<br>"Synovial sarcoma",<br>" Blastoma " |
| Thoracic and respiratory | "Adenocarcinoma of the lung",<br>"Basaloid squamous cell lung carcinoma",<br>"Bronchial adenomas/carcinoids",<br>"Giant-cell carcinoma of the lung",<br>"Large-cell lung carcinoma",<br>"Large cell lung carcinoma with rhabdoid phenotype",<br>"Laryngeal cancer",<br>"Mesothelioma",<br>"Non-small cell lung cancer",<br>"Non-small cell lung carcinoma",<br>"Pleuropulmonary blastoma",<br>"Sarcomatoid carcinoma of the lung",<br>"Small cell lung cancer",<br>"Squamous-cell carcinoma of the lung",<br>"Thymoma and thymic carcinoma" |

| | |
|---|---|
| HIV/AIDS related | "AIDS-related cancers",<br>"Kaposi sarcoma" |

**Table S8. Regular expressions for Cancer Types.** The list was derived from Wikipedia[3].

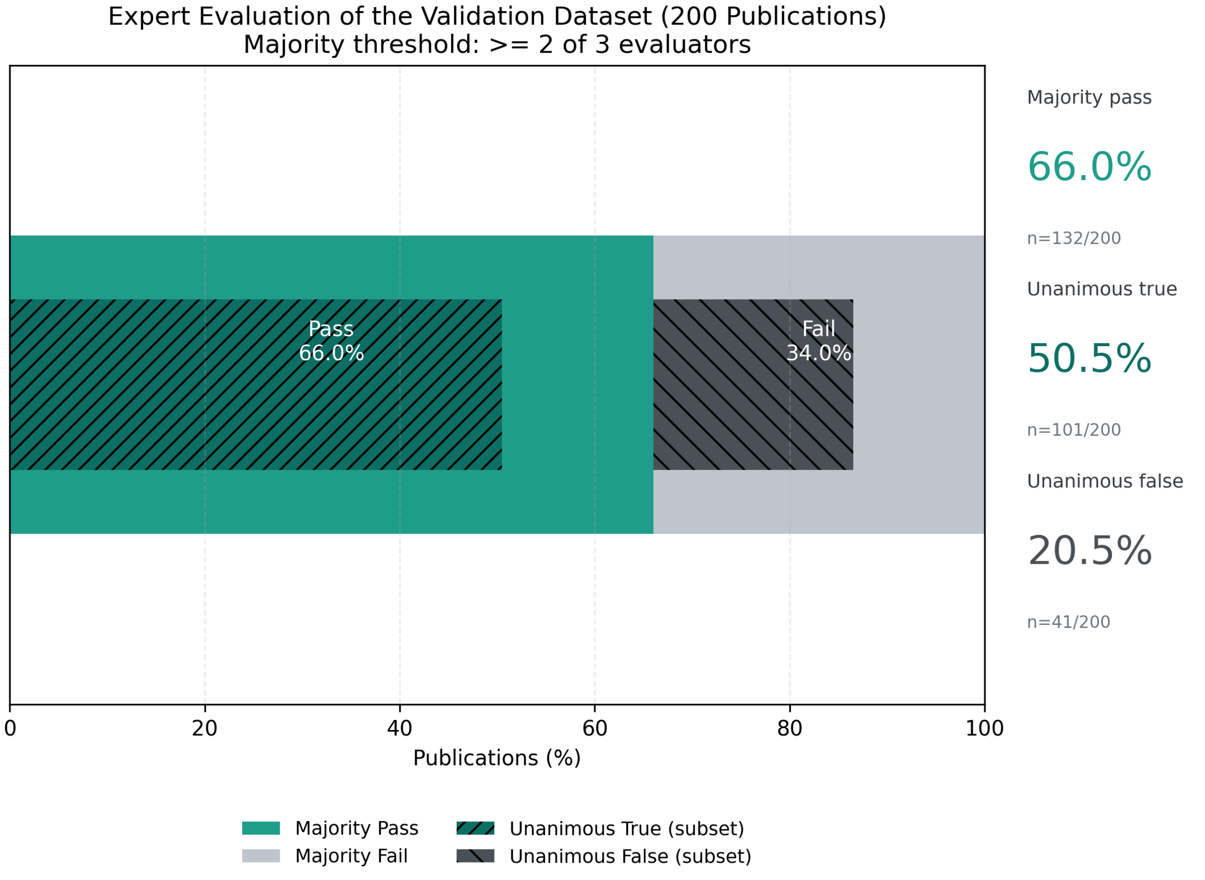


**Figure S1.** Results of the human expert curation on the 200 publications contained in the validation dataset. The same dataset is used to estimate the number of false positives in the keyword-based dataset. Majority Pass and Fail represent the percentage of documents that have been deemed relevant or irrelevant by the majority of experts (at least 2/3). Unanimous True or False represent the percentage of documents considered relevant or irrelevant by all the experts.

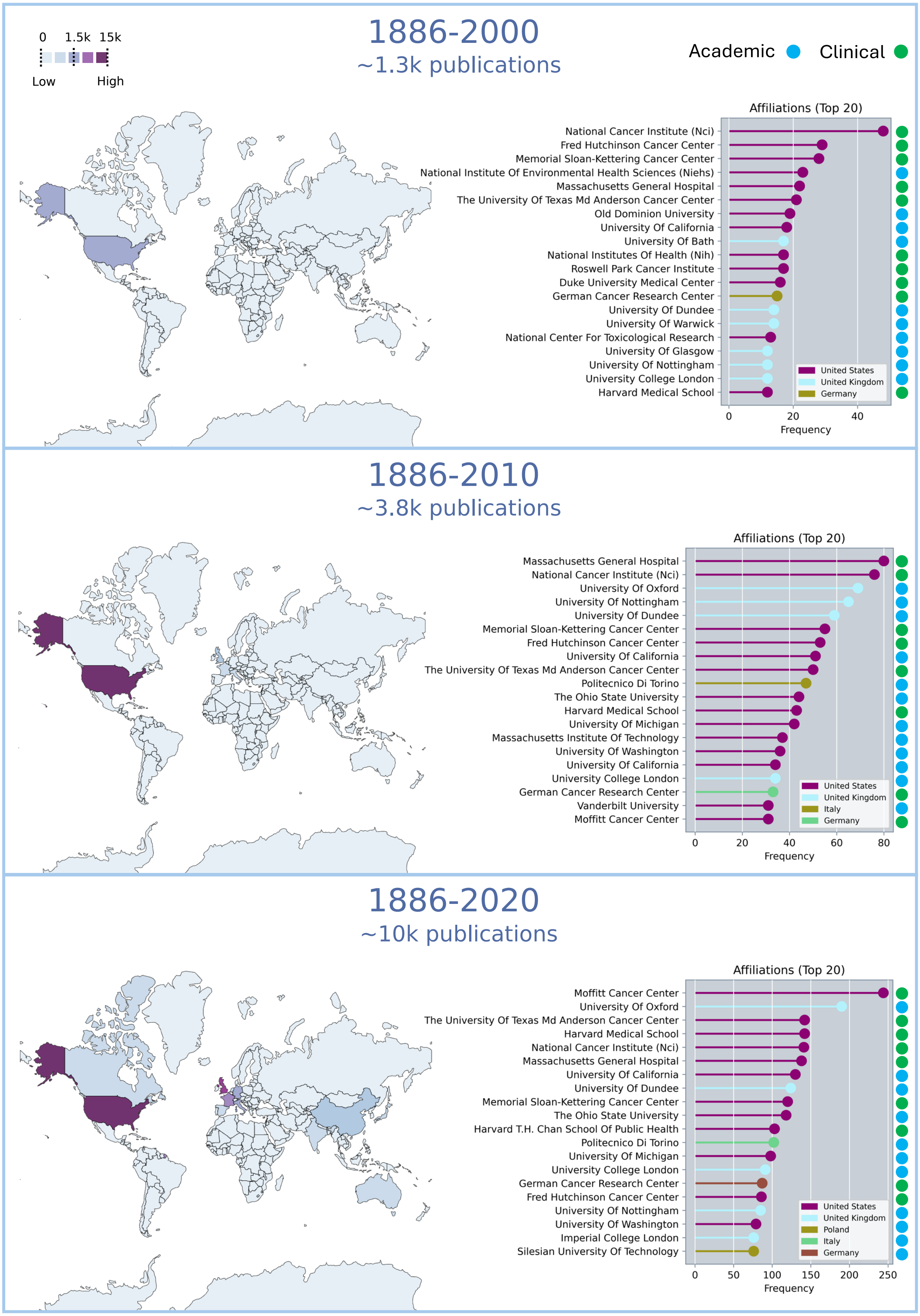


**Figure S2. Geographic distribution of mathematical research in oncology over time (LLM-curated dataset).**
To characterize the temporal evolution of geographic contributions, the LLM-based dataset was partitioned into four subsets, each corresponding to a cumulative increment of publications (1886–2000; 1886–2010; 886–2020; 1886–2025). The full dataset (1886–2025) is shown in **Figure 2** of the main text, while the three preceding subsets are displayed here (top: until 2000; middle: until 2010; bottom: until 2020). Across time frames, the distribution consistently reflects the prominence of Western institutions, particularly those in the U.S. and the U.K., with recent growth in contributions from other countries. Notably, each of the analyzed time frame see a strong representation of both clinical and academic institutions, demonstrating the inherently translational nature of mathematical research in oncology.

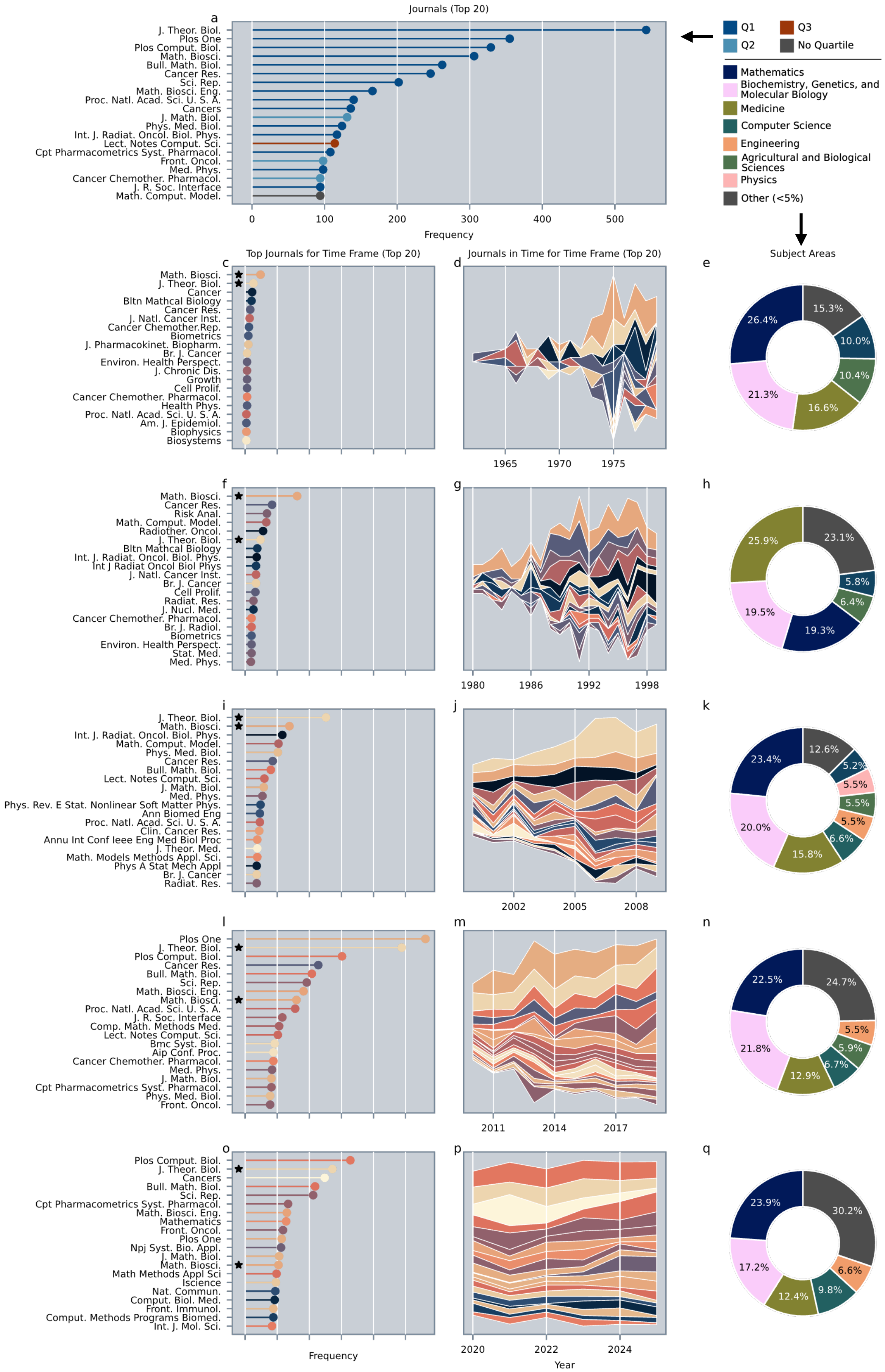


**Figure S3. Most represented journals and subject areas in the LLM-curated dataset over time.** a) Most represented journals in the LLM-curated dataset (top 20), colored by quartile. Most of the listed journals belong to the fields of Computational and Mathematical Biology (e.g., *The Journal of Theoretical Biology*, *The Bulletin of Mathematical Biology*, *PLoS Computational Biology*), but four multidisciplinary (*PLoS ONE*, *Scientific Reports*, *PNAS*, *J. R. Soc. Interface*) and four oncological journals (*Cancer Research*, *Cancers (Basel)*, *Cancer Chemotherapy and Pharmacology*, *Frontiers in Oncology*) are also present in the ranking. To characterize the temporal evolution of journals and scientific areas, we partitioned the LLM-curated dataset into 5 time frames: 1886-1980 (b-d), 1980-200 (e-g), 2000-2010 (h-j), 2010-2020 (k-m), 2020-2025 (n-p). In the left column, we show the top 20 journals for the four different time frames. We noted with a black star the journals that are present throughout all the rankings. In the central column, we show the number of publications for each journal over time in the selected span. In the right column, we show the distribution of journal subject areas. The group "Other" includes all ASJC areas representing less than 5% of the publications in the query-based dataset.

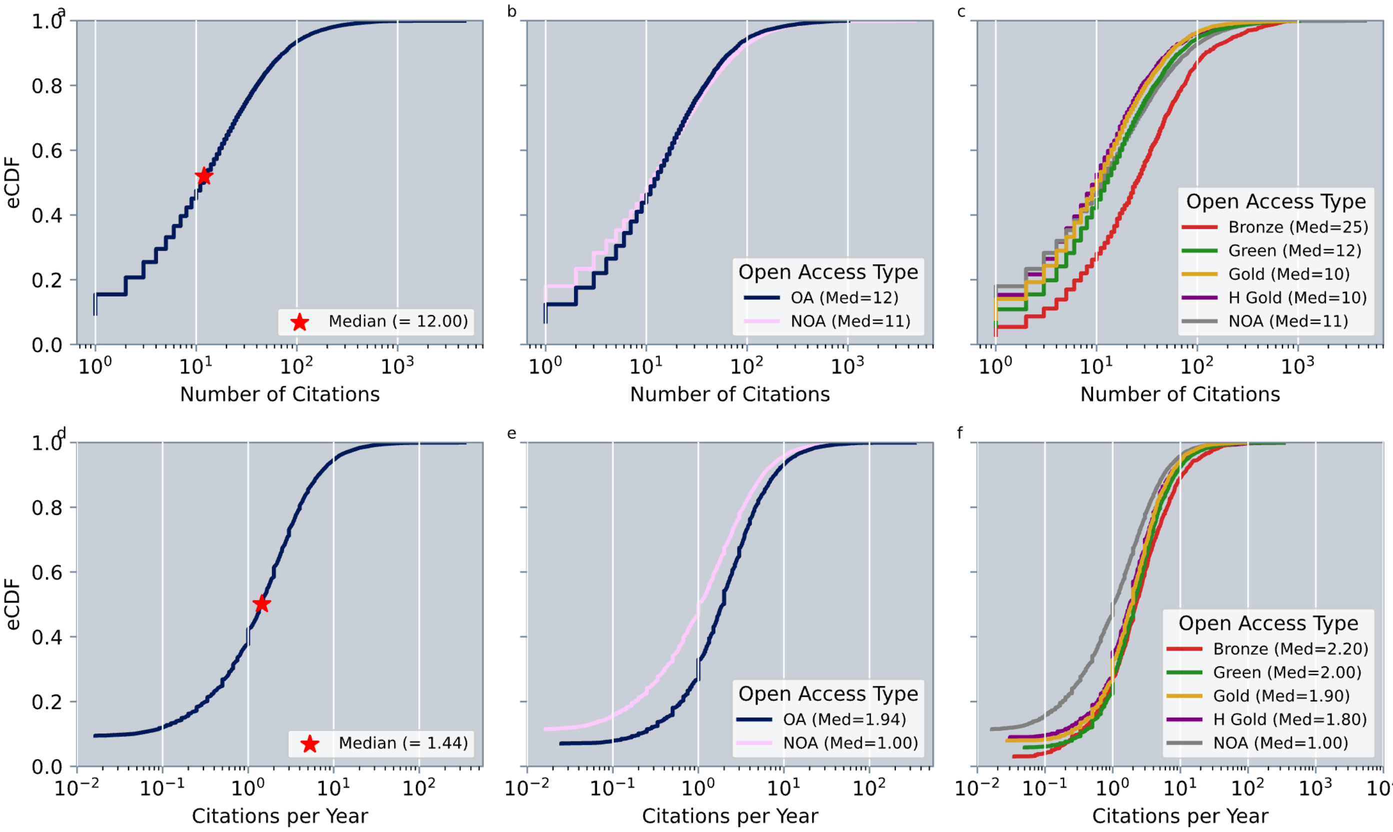


**Figure S4. Citation patterns in the query-based dataset. a)** Empirical cumulative distribution (eCDF) of citation counts; the star marks the median. **b)** eCDF of citation counts stratified by Open Access (OA, n=6217) versus Non-OA (NOA, n=7561); median values are reported in parentheses. **c)** eCDF of citation counts stratified by OA type (Bronze, Green, Gold, Hybrid Gold), with medians in parentheses. Number of documents for each category: Bronze (n=1183), Green (n=4799), Hybrid Gold (n=1113), Gold (n=3632). **d–f)** eCDFs of citations per year. Statistical comparisons between distributions are provided in **Table S1**.

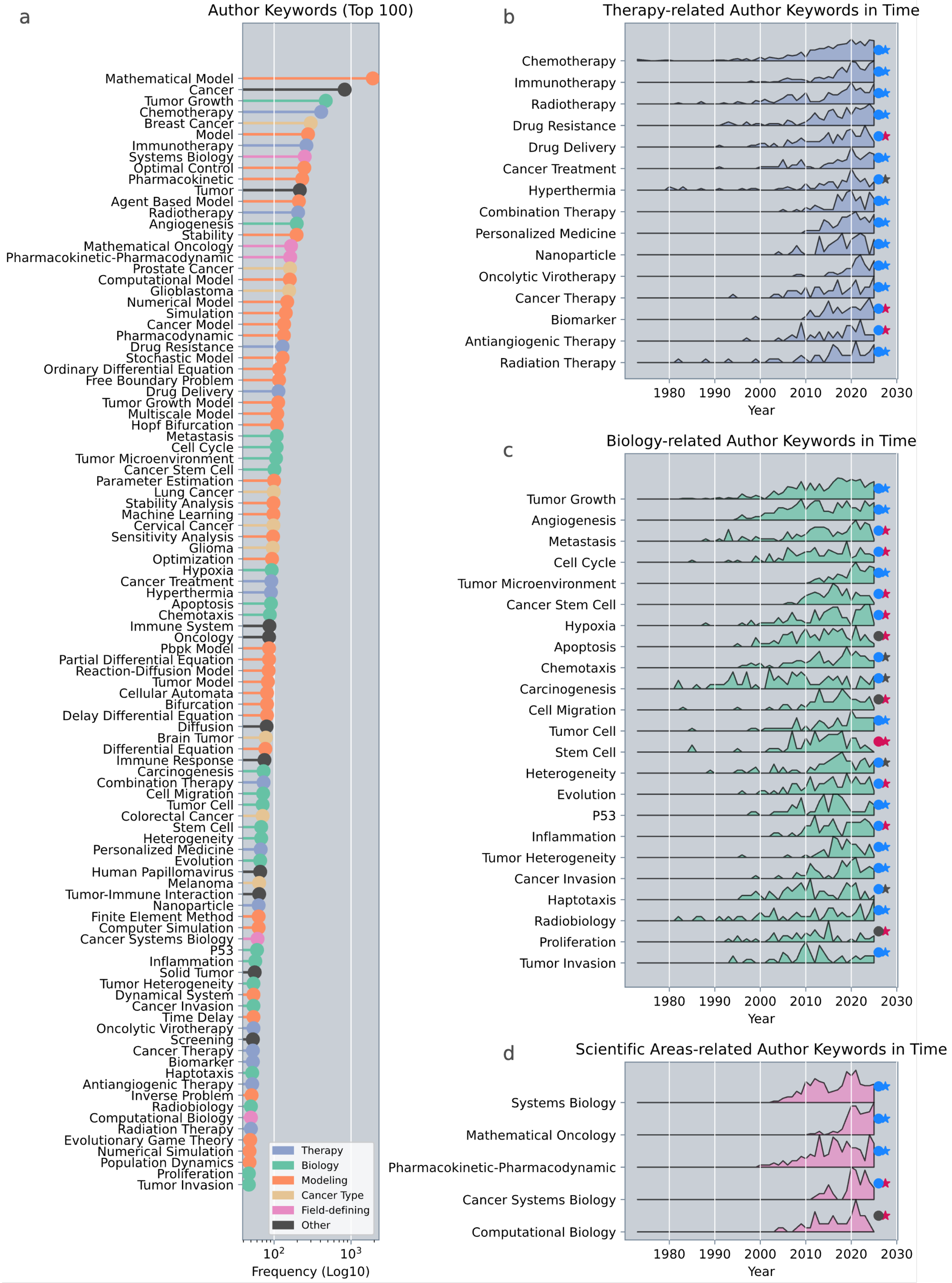


**Figure S5. Most represented AKs in the llm-curated dataset.**
**a)** Top 100 AKs in the dataset, colored by category. Usage in time of the AKs associated to **b)** therapeutic modalities, **c)** cancer biology, and **d)** scientific areas. Each temporal trend is normalized independently for each AK. The average increase or decrease of each AK is indicated by a circle or a star on the right side of the curve. A blue circle denotes an average increase in term usage over the past 30 years (1995–2025), a red circle denotes a decrease, and a grey circle denotes no change. Similarly, a blue star indicates an average increase over the past 10 years (2015–2025), and a red star indicates a decrease, and grey indicates no change.

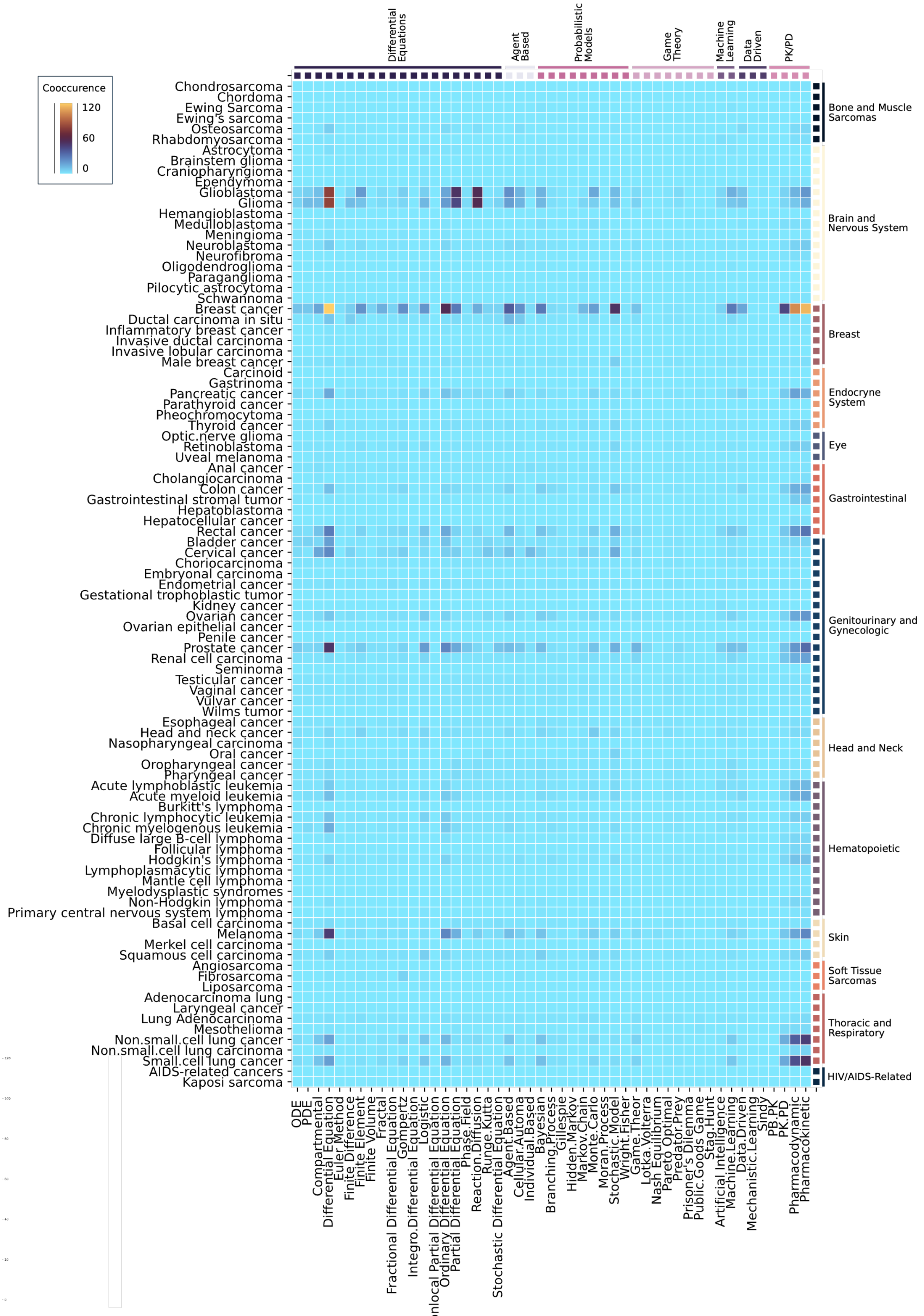


**Figure S6. Co-occurence matrix between regex associated with cancer types (y labels) and modeling methods (x labels).** The category of each regex is annotated on the opposite side. Consistent with the historical prominence of differential equations, PK/PD frameworks, and probabilistic modeling, these approaches dominate across most cancer types studied in mathematical research in oncology. A notable exception is ductal carcinoma in situ, which has been examined predominantly using agent-based models. Highly investigated tumors (e.g., breast cancer, prostate cancer, glioma, glioblastoma) tend to be associated with multiple modeling approaches, suggesting that tumor complexity rather than methodological preference drives the application of diverse mathematical techniques.

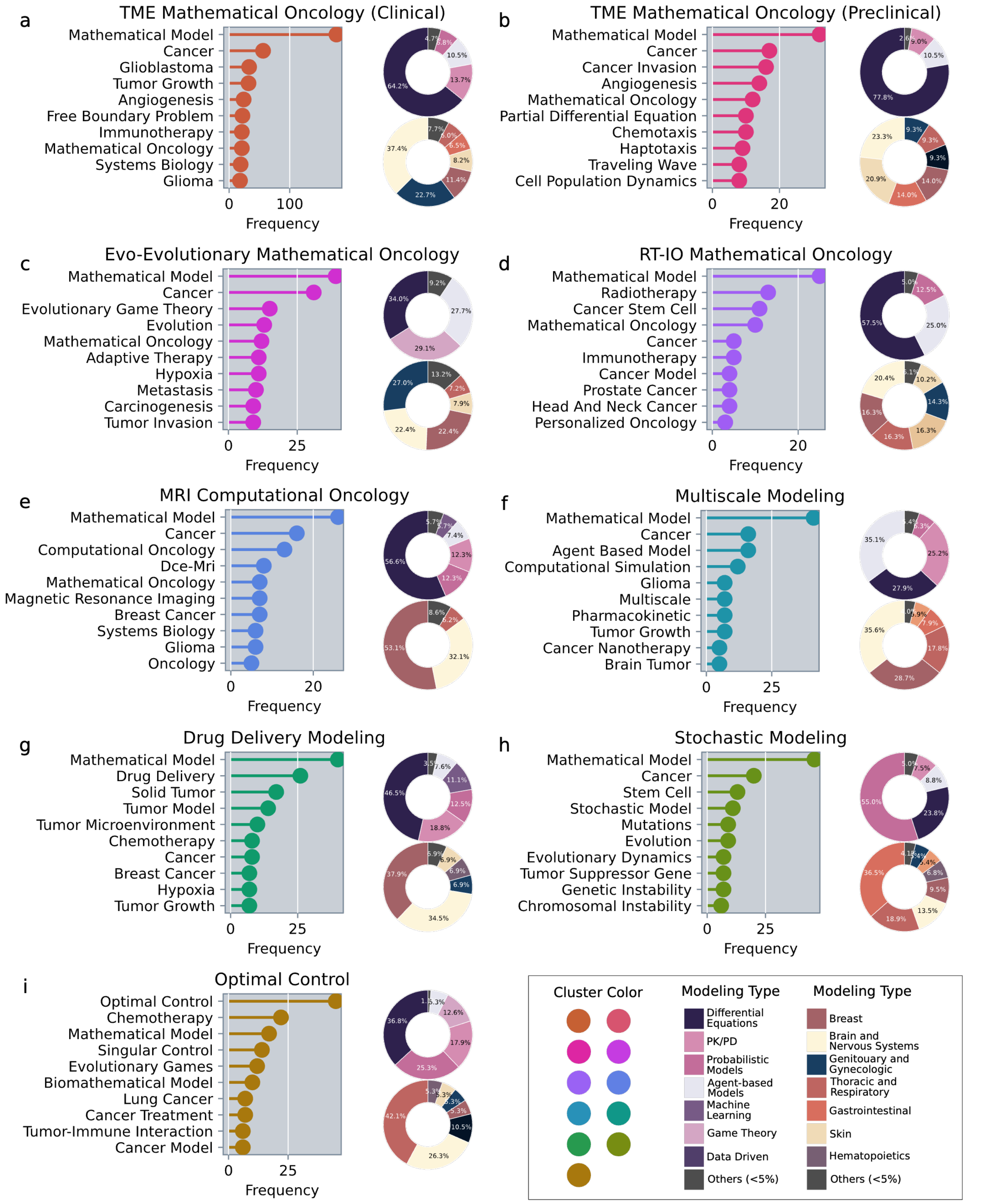


**Figure S7. Most prominent themes across the nine author clusters. a-i)** each panel shows the 10 most frequent keywords, the frequency of mathematical modeling techniques, and the prevalence of cancer types in each cluster. The color of each lollipop plot correspond to the cluster color in **Fig. 5** (the name of each cluster is also reported above each subpanel). The legend represent the color associated with each modeling class, and cancer type.

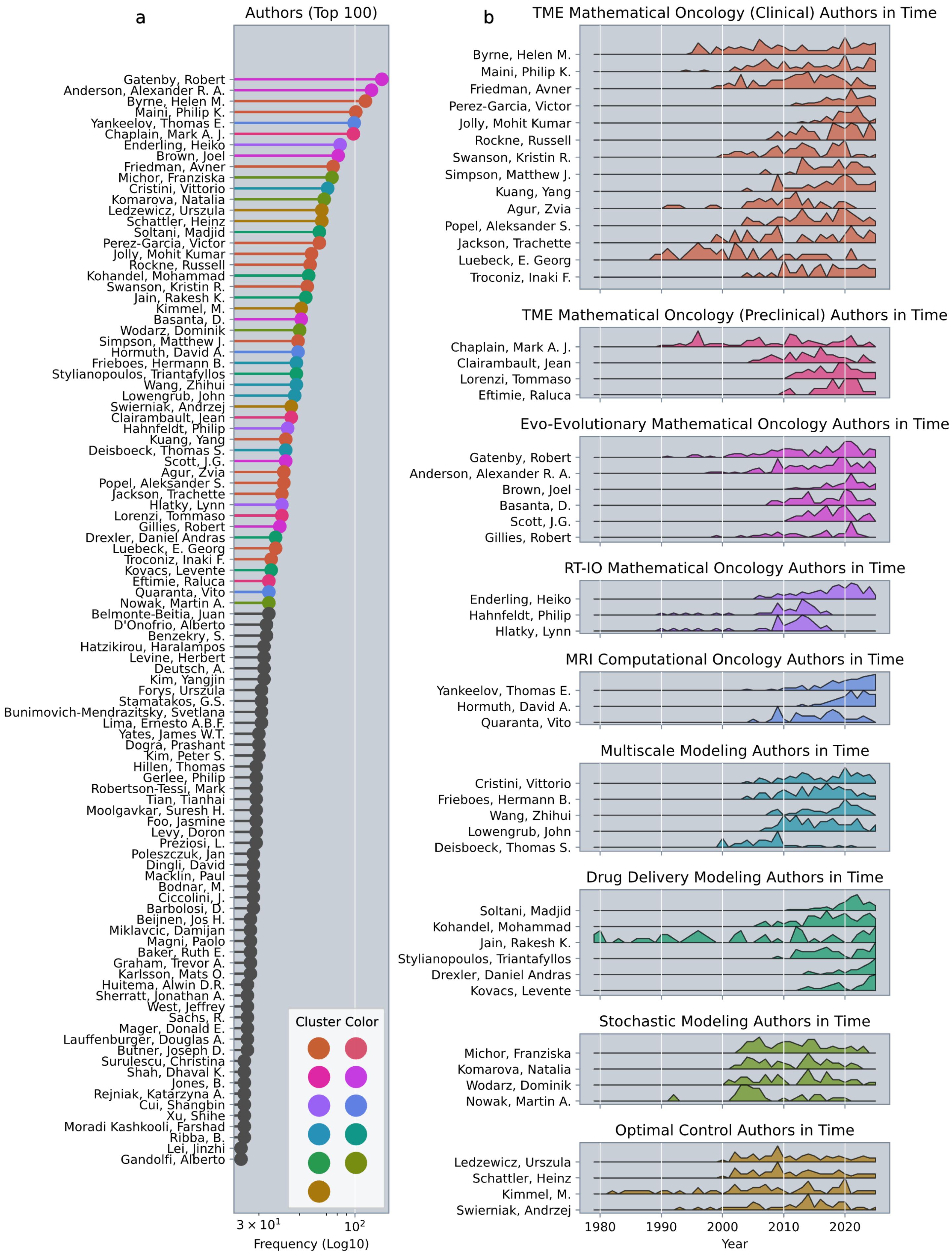


**Figure S8. Top 100 authors and their temporal trends in the LLM-curated dataset. a)** top 100 authors in the LLM-curated dataset. Note that the number of documents in the dataset does not reflect the total productivity of each authors, but rather the number of document that our pipeline selected as relevant for mathematical modeling in oncology. **b)** Number of documents in time for each author, independently normalized for each one.

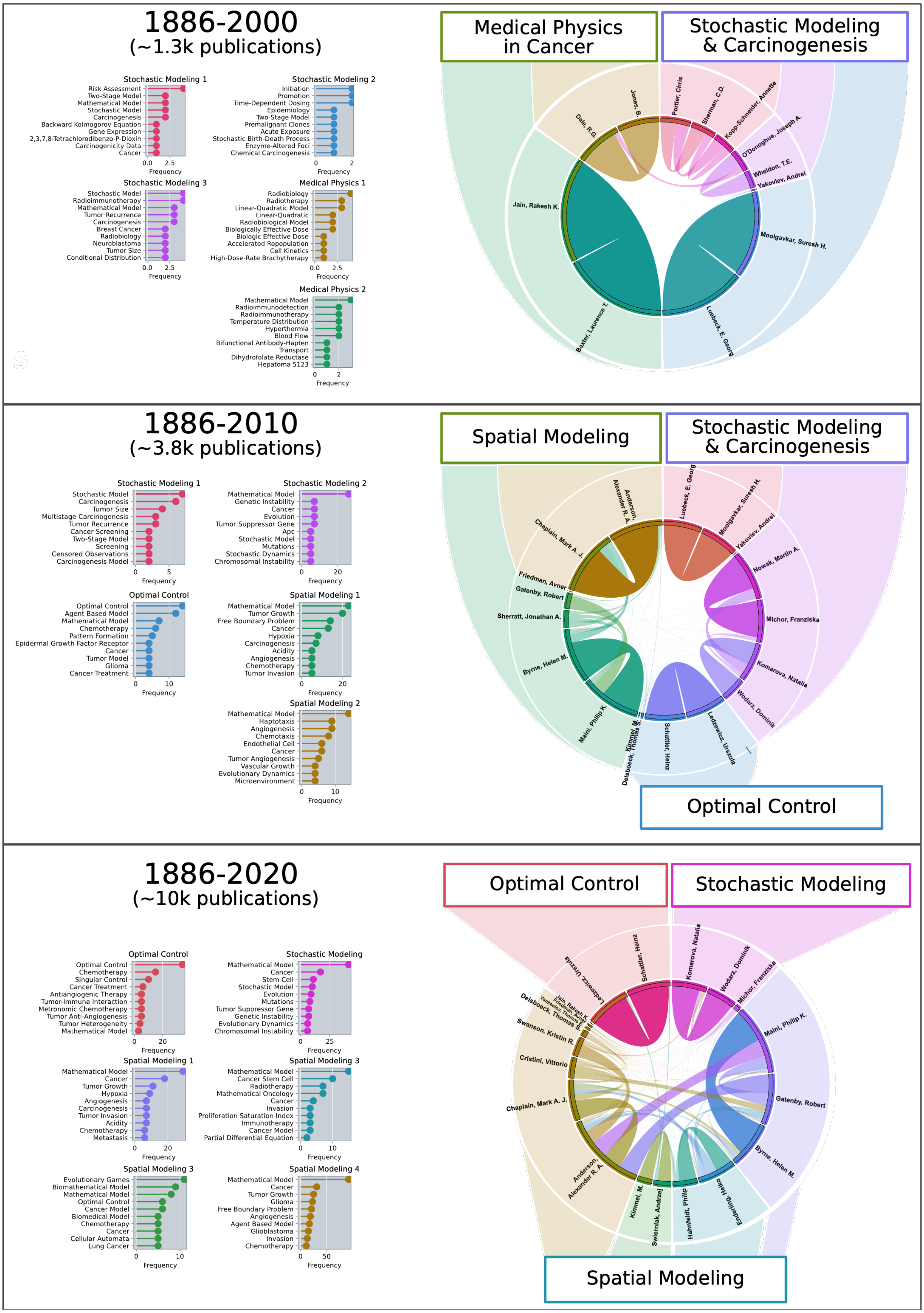


**Figure S9. The emergence of spatial modeling in mathematical research in oncology.** The current scientific landscape evidences a broad and shared interest in spatial modeling applied to cancer and to the tumor microenvironment (Fig. 5). The bibliographic coupling of the top 20 authors in time reveals that this interest emerged only in the last 25 years. In the 20th century, the major applications of mathematical research to oncology were limited to statistical modeling (especially to understand carcinogenesis), and medical physics (e.g. radiobiology). With the advent of the 21st century, and the understanding of cancer as a living ecosystem, spatial modeling groups emerged prominently, until the formation of modern Mathematical oncology.
